# KIR Expression Defines a Transcriptionally Distinct CD8+ T-cell Population that Confounds Antigen-Specific T-cell Detection

**DOI:** 10.64898/2026.07.18.739377

**Authors:** Andreas H. Kongsgaard, Janine Kemming, Sofie Ramskov, Annie Borch, Yogesh Basavaraju, Alberte Obitz Mogensen, Tsvetanka Rangelova, Mohammad Kadivar, Sine Reker Hadrup, Sunil Kumar Saini

## Abstract

HLA-C–restricted CD8^+^ T-cell responses remain poorly defined compared with HLA-A and HLA-B responses, despite the central role of HLA-C as both a peptide-presenting molecule and a ligand for killer-cell immunoglobulin-like receptors (KIRs). This dual function creates a major challenge for pHLA multimer-based antigen-specific T-cell analysis, as HLA-C multimers may bind CD8^+^ T-cells through either antigen-specific TCRs or KIRs.

Here, we show that KIR expression on CD8^+^ T-cells drives TCR-independent binding to pHLA-C multimers, leading to overestimation of HLA-C–restricted antigen-specific T-cell responses. Using HLA-mismatched donor settings and cancer neoantigen pHLA-C multimers, we demonstrate that apparent multimer-positive CD8^+^ T-cells can arise from KIR-mediated recognition rather than cognate TCR specificity. We further show that KIR blockade before pHLA-C multimer staining removes non–TCR-specific binding, and applying this strategy to large-scale analysis of CMV, EBV, and SARS-CoV-2 antigens, we identify TCR-specific HLA-C–restricted CD8^+^ T-cell populations. Importantly, KIR-mediated pHLA-C binding is peptide-selective rather than uniform across all peptide–HLA-C complexes. Single-cell transcriptomic, phenotypic, and clonotypic analyses further demonstrate that KIR-mediated and TCR-mediated pHLA-C multimer binding define transcriptionally and clonally distinct CD8^+^ T-cell populations.

Together, these findings establish KIR-mediated pHLA-C recognition as a major confounder in HLA-C multimer-based T-cell analysis. By separating KIR- from TCR-mediated binding, our approach provides a practical framework for accurate discovery and characterization of HLA-C–restricted antigen-specific CD8^+^ T-cells across viral infection and cancer.

## Introduction

T-cells play a key role in the adaptive immune response with the T-cell receptor (TCR) of CD8^+^ T-cells recognizing class I major histocompatibility complex (MHC-I) associated peptides presented by all nucleated cells. Understanding CD8^+^ T-cell recognition and functional response is vital for improving therapies for viral infections, autoimmune diseases, and cancer. Current techniques to explore peptide-MHC (pMHC) interactions include the use of fluorescent dyes, molecular barcodes, or metal tags, allowing parallel analysis of multiple peptide specificities (1–4). Identifying antigen-specific T-cells facilitates precise targeting of tumors and viral infections through adoptive cell therapy while contributing to our understanding of immune responses in health and disease.

MHC-I molecules are categorized into human leukocyte antigen (HLA)-A, -B, and -C in humans. While both HLA-A and -B have been widely studied for their interaction with T-cells, much less is known about HLA-C. This is likely due to the perception of HLA-C as a minor player in T-cell-mediated immunity, given that its surface expression is significantly lower than HLA-A and -B, reported up to 13–18 fold (5–7). Interestingly, while HLA-A and -B are often downregulated in virus-infected cells, HLA-C expression is relatively preserved (6). HLA-C-restricted T-cells have been implicated in diverse disease settings. Higher HLA-C expression has been associated with enhanced T-cell responses and slower disease progression in HIV-1 infection (8). In contrast, elevated HLA-C expression and specific HLA-C alleles have also been linked to immune-mediated diseases, including Crohn’s disease and psoriasis (6,9,10). Additionally, HLA-C-restricted CD8^+^ T-cells have been shown to play a role in various cancers (11–13). More recently, KIR^+^CD8^+^ T-cells have been proposed to represent a regulatory T-cell population, including circulating tumor-reactive KIR^+^CD8^+^ T-cells implicated in the suppression of anti-tumor immunity in melanoma (14,15). Together, this emphasizes the importance of studying HLA-C-restricted antigens due to their critical involvement in various disease settings.

While CD8^+^ T-cell binding to HLA-A and -B is primarily driven by TCR-mediated interactions, HLA-C binding can frequently occur through KIR-mediated binding, as well as TCR-mediated binding, and differentiation between the two binding options is challenging using pHLA multimers, when KIRs are expressed on CD8^+^ T-cell populations. Unlike the TCR, KIR’s interaction with HLA-C is based less on peptide specificity and more on binding to conserved sequences of the HLA-C molecule (16,17). HLA-C allotypes are classified into C1 and C2 groups based on amino acid residue 80, with Asn80 defining C1 and Lys80 defining C2 (18). Most HLA-C–binding KIRs preferentially recognize one of these two allotype groups, with additional peptide-dependent modulation of binding reported for specific KIR–HLA-C interactions (16,19–21). With approximately 5% of CD8^+^ T-cells reported to express KIRs (22), the dual recognition by TCRs and KIRs poses challenges for accurately identifying TCR-driven HLA-C restricted interactions and constraining the ability to improve therapies through targeting antigens presented via this allele. This study demonstrates that pHLA-C multimers, carrying virus-derived peptides or neoepitopes (derived from sequences with cancer-specific mutations), showed extensive KIR-driven binding in healthy donors as well as in cancer patient samples, thus limiting identification of antigen-specific T-cells with TCRs recognizing HLA-C presented epitopes. By blocking KIR receptors using KIR-specific antibodies, we developed a strategy to differentiate KIR-and TCR-driven binding to pHLA-C, enabling large-scale T-cell detection approach for reliable identification of HLA-C-restricted antigen-specific T-cells.

## Results

### Expression of KIRs on CD8^+^ T-cells hinders the discovery and analysis of antigen-specific T-cells

pHLA multimers are widely used to identify and evaluate antigen-specific T-cells. Epitope-specific pHLA multimers identify T-cells by binding to their cognate TCRs. However, in the context of HLA-C, pHLA multimers can also bind to the KIR molecules, expressed by a subset of CD8^+^ T-cells (22,23). Such KIR-mediated binding of pHLA multimers can lead to a false discovery of antigen-specific T-cells (Fig. 1A). KIR expression was detected on 2.5% (range 0.32-16.7%; n=41) of CD8^+^ T-cells from healthy donors, compared with 27.6% on NK cells (range 11-59%; n=21) using a KIR2DL1/DS1/DS3/DS5 antibody (Fig. 1B). While KIR-driven pHLA multimer binding to NK cells can easily be distinguished by NK linage markers, we sat out to explore if CD8^+^ T-cell binding of pHLA-C multimers can be separated into TCR- and KIR-driven interactions. To test this, we evaluated the binding of HLA-C*0202 pHLA tetramers, holding a neoantigen peptide (VAALATSFL), previously identified from somatic mutational profiling in a cancer patient. Despite the cancer- and patient-specific nature of the peptide, these pHLA-C multimers showed binding to CD8^+^ T-cells across a range of healthy donors, expressing both matched (C*02:02) and mismatched (C*05:01, C*07:02) HLA-C molecules, respectively (Fig. 1C). However, such CD8^+^ T-cells didn’t show any functional activation (no cytokine release) upon stimulation with the peptide, suggesting a TCR-independent binding of HLA-C-specific multimers (Fig. 1C).

**Figure 1:**
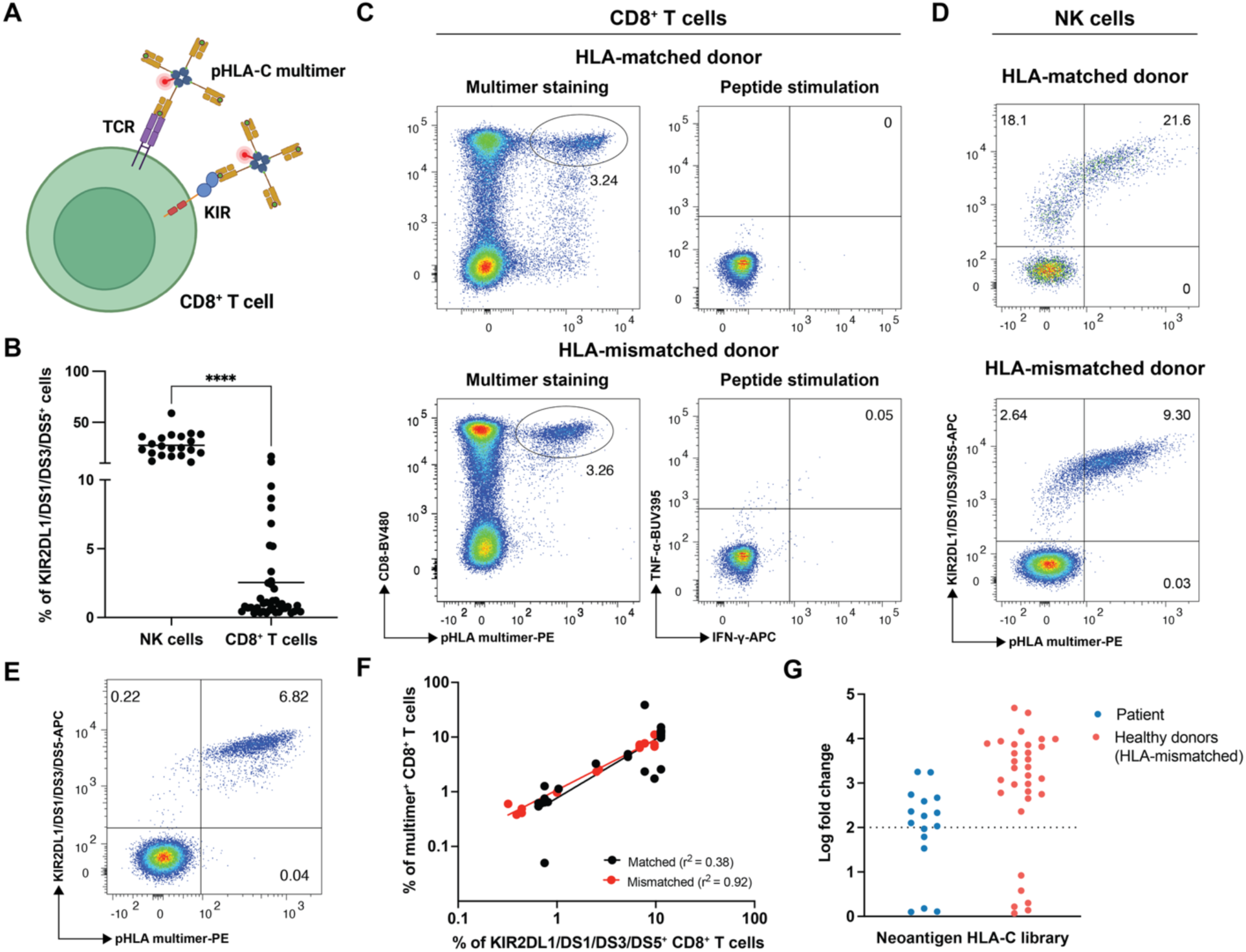
KIR expression on CD8^+^ T-cells confounds pHLA-C multimer-based identification of antigen-specific T-cells. **A)** Representation of pHLA-C multimer binding to CD8^+^ T-cells via TCR or KIR. **B)** KIR2DL1/DS1/DS3/DS5 expression on NK cells and CD8^+^ T-cells in healthy donor PBMCs. Dots represent individual donors and horizontal lines indicate the mean. Mean expression was 27.6% on NK cells (range, 11-59%, n=21) and 2.5% on CD8^+^ T-cells (range, 0.32-16.7%, n=41). Statistical significance was assessed using a two-sided Wilcoxon matched-pairs signed-rank test on donors with paired NK-cell and CD8^+^ T-cell measurements (n=21, p<0.0001). **C)** Flow cytometry plots showing pHLA-C multimer staining (C*02:02-VAALATSFL) and peptide-induced IFN-γ and TNF-α production by CD8^+^ T-cells from an HLA-C*02:02-matched donor and a non-HLA-C*02:02 donor (HLA-mismatched). Multimer staining was tested in HLA-C*02:02-matched donors (n=4) and a non-HLA-C*02:02 donor (n=1), while peptide-induced cytokine production was assessed only in the donors shown. **D)** Co-staining of NK cells with C*02:02-VAALATSFL multimer and KIR2DL1/DS1/DS3/DS5 in HLA-C allotype-matched and mismatched donors. **E)** Representative C*02:02-VAALATSFL multimer staining relative to KIR2DL1/DS1/DS3/DS5 expression in CD8^+^ T-cells from an HLA-C*02:02-mismatched donor. **F)** Correlation between pHLA-C multimer binding and KIR2DL1/DS1/DS3/DS5 expression across HLA-C allotype-matched donors (n=47) and mismatched donors (n=17). **G)** Analysis of a MDS patient-specific neoantigen library restricted to HLA-C*02:02 (n=16) to identify CD8^+^ T-cell binding using DNA-barcoded pHLA multimers and compared with an HLA-C*02:02-mismatched healthy donor. Significant pHLA binders are shown above the dotted line (Log2 fold change >2 and p<0.001).

The multimer binding observed in a non-HLA restricted fashion and the lack of peptide-responsiveness hinted to a KIR-mediated binding of these multimers. To validate this, we next evaluated if these multimers also recognized NK-cells, and if the frequency of KIR-expressing NK-cells correlates with that of pHLA multimer-binding cells. Indeed, NK-cells of the two donors showed binding to the pHLA multimers consistent with their frequency of KIR expression (Fig. 1D). Finally, we show that all multimer-binding CD8^+^ T-cells and NK-cells co-expressed KIR2DL1/DS1/DS3/DS5, suggesting that KIRs are mediating pHLA-C binding in a TCR-independent manner. (Fig. 1E). Furthermore, pHLA-C multimer binding to CD8^+^ T-cells correlated strongly with the KIR2DL1/DS1/DS3/DS5 expression in both HLA-mismatched (n=11; r²=0.92, p<0.0001) and HLA-matched (n=7; r²=0.38, p<0.0001) donors (Fig. 1F). Altogether, our data suggests the KIR-molecules are implicated in pHLA-C-multimer binding to CD8^+^ T-cells, and that such binding results in staining intensities very similar to that of ‘true’ TCR-driven pHLA binding, albeit with often larger T-cell population sizes.

To investigate whether KIR-mediated binding could bias pHLA multimer-based detection of antigen-specific T-cells, we examined published large-scale datasets reporting antigen-specific CD8 T-cells (24–27), and found that HLA-C-specific T-cell populations were significantly overrepresented compared to HLA-A and -B-restricted responses (Supplementary Fig. 3A, Supplementary Table 1). This is despite the significantly lower expression of HLA-C at transcriptomic and protein levels, and the lower cell surface peptidome compared to HLA-A and HLA-B molecules (Supplementary Fig. 3B-E). Thus, such disproportional identification of HLA-C-binding T-cells could be explained by binding of pHLA-C multimers to the subset of CD8^+^ T-cells expressing KIRs instead of TCRs. To examine this further, we evaluated T-cell binding using pHLA-C multimers loaded with 16 neopeptides derived from an MDS patient, predicted to bind HLA-C*02:02 molecules. Using DNA-barcoded pHLA multimers (4), T-cell binding was measured in the patient sample and two healthy donors with HLA-C*02:02 mismatch background. For comparison, we additionally stained the samples using 49 HLA-A and 38 HLA-B-multimers also presenting neopeptides (Supplementary Fig. 4, Supplementary Table 2). The majority of pHLA-C multimers showed significant binding (LogFC>2, p<0.001) to CD8^+^ T-cells both in MDS patients and HLA-mismatched healthy donors (Fig. 1G), suggesting a binding receptor that is both HLA and peptide agnostic. In contrast, none of the HLA-A and -B restricted multimers displayed T-cell binding (Supplementary Fig. 4). Thus, providing strong evidence of KIR-mediated interference in large-scale antigen-specific T-cells discoveries.

### KIR-blocking differentiates KIR- and TCR-driven binding of HLA-C multimers

Next, we tested if pre-incubation with an anti-KIR antibody could block KIR-mediated interactions with pHLA multimers, thereby enabling the identification of TCR-specific pHLA binding (Fig. 2A). We incubated PBMCs with an anti-KIR2DL1/DS1/DS3/DS5 antibody before the addition of HLA-C-multimers. This approach completely abolished pHLA-C multimer binding to both CD8^+^ T-cells and NK cells in an HLA-mismatched context (Supplementary Fig. 5, Fig. 2B, left), while non-HLA-C multimers such as the HLA-A*02:01/CMV-NLV multimer retained CD8^+^ T-cells binding, even the presence of KIR blocking antibodies (Fig. 2B, Right).

**Figure 2:**
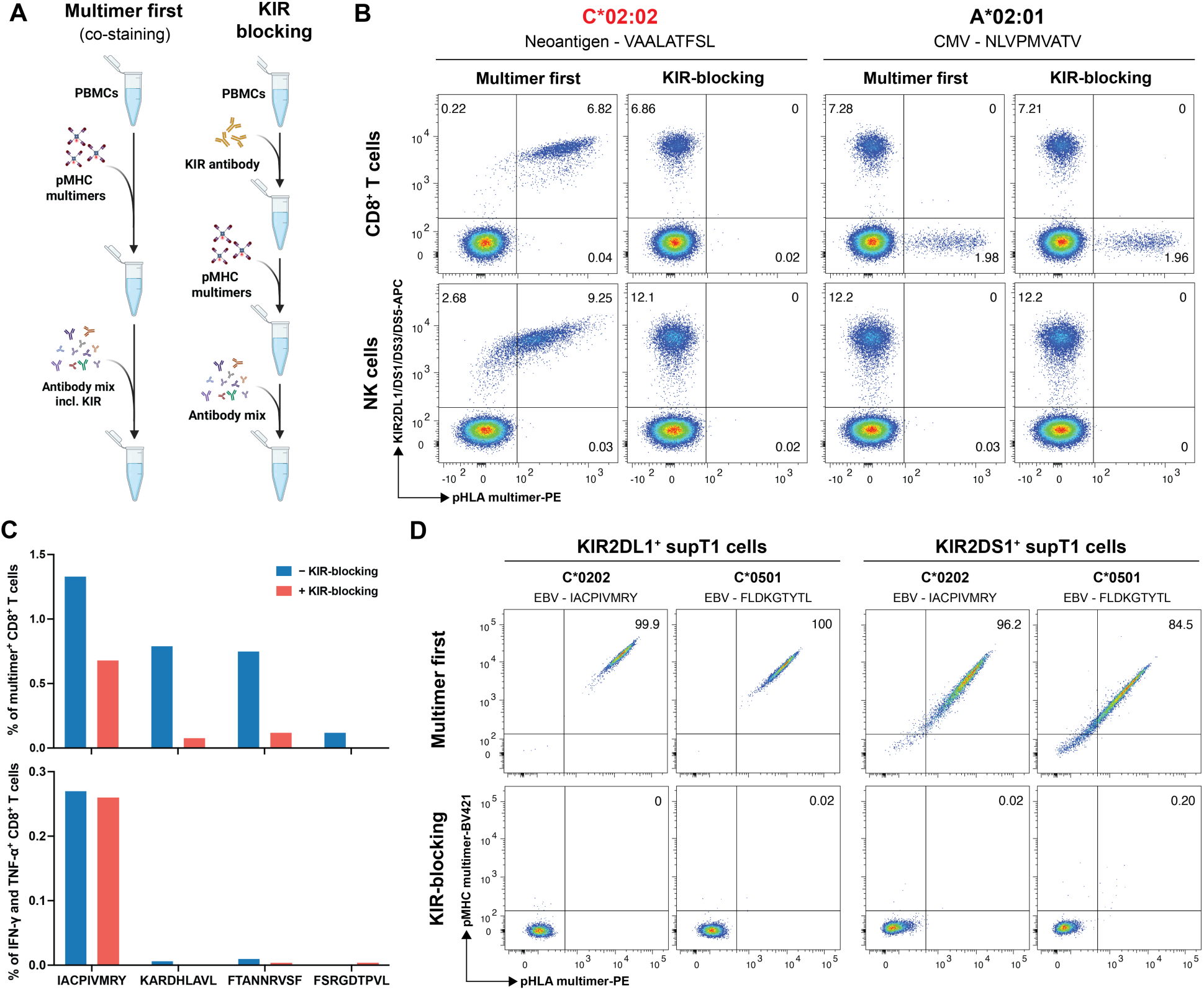
KIR-blocking separates KIR- and TCR-mediated pHLA-C binding. **A)** Schematic of anti-KIR antibody-mediated KIR-blocking strategy for pHLA-C multimer staining for antigen-specific CD8^+^ T-cell analysis. **B)** Left, flow cytometry plots showing pHLA-C multimer (C*02:02-VAALATFSL) staining of CD8^+^ T-cells and NK cells using KIR2DL1/DS1/DS3/DS5 as a co-stain, added after multimer staining, or as a KIR-blocking antibody in an HLA-mismatched donor. Right, similar analysis using pHLA multimers carrying the HLA-A*02:01-restricted CMV epitope NLVPMVATV in healthy donor PBMCs. **C)** Top, Frequency of CD8^+^ T-cells binding to HLA-C-restricted CMV peptides with and without KIR-blocking identified using combinatorial tetramer analysis in healthy donor PBMCs (Supplementary Fig. 6). Bottom, TCR-mediated peptide-induced IFN-γ and TNF-α secretion by CD8^+^ T-cells by the identified peptides with or without KIR-blocking (Supplementary Fig. 7). **D)** Flow cytometry plots showing pHLA-C multimer binding to SUP-T1 cells transduced with KIR2DL1 or KIR2DS1 under KIR co-staining (multimer-first staining) and KIR-blocking conditions.

To distinguish TCR-mediated recognition of pHLA-C from KIR-driven multimer binding, we incorporated KIR blocking into the pHLA-C multimer analysis workflow. We assessed CD8^+^ T-cell recognition of CMV- and EBV-derived peptides presented by HLA-C using a library of 46 peptides predicted to bind HLA-C*02:02 (Supplementary Table 3) and tested the resulting multimers for T-cell binding in PBMCs from two HLA-C*02:02-matched healthy donors. T-cell reactivity was assessed via combinatorial fluorochrome labeling of pHLA multimers representing different specificities (1), performed in parallel with and without KIR blockade. In the absence of KIR-blocking, four pHLA-C multimer specificities showed binding to CD8^+^ T-cells (0.12-1.33% of CD8).

However, only one of these multimers retained substantial binding in the KIR-blocking condition, indicating TCR-dependent binding, whereas the remaining three specificities showed minimal or no binding (Supplementary Fig. 6). All four peptide specificities were further tested for functional activation of CD8^+^ T-cells upon stimulation. Only IACPIVMRY peptide, that had retained substantial binding in the KIR-blocking setup, induced cytokine (IFN-γ and TNF-α) secretion, indicating a TCR-specific binding of HLA-C*02:02-IACPIVMRY multimers, while the remaining peptides revealed binding through KIRs, and such T-cells do not respond to peptide stimulation (Fig. 2C, Supplementary Fig. 7). Importantly, KIR-blocking did not impair TCR signaling, as IFN-γ and TNF-α secretion remained comparable in the presence and absence of KIR-blocking (Fig. 2C, Supplementary Fig. 7). Furthermore, under KIR-blocking conditions, only 50% of the IACPIVMRY-specific pHLA multimer binding was retained compared to the non-blocking condition. This suggests that even when binding is TCR-specific, the presence of KIR-expressing CD8^+^ T-cells can significantly affect the apparent frequency of antigen-specific CD8^+^ T-cells in multimer-based assays. The peptide, IACPIVMRY, derived from EBV, has not previously been reported as immunogenic in the context of HLA-C*02:02 (IEDB), highlighting it as a novel HLA-C-restricted T-cell epitope.

As the C2 group of KIR ligands include several HLA-C molecules (28), we tested if KIR-blocking using the anti-KIR2DL1/DS1/DS3/DS5 antibody is also effective against other C2 group HLAs such as HLA-C*05:01. SupT1 cell lines transduced to express KIR2DL1 or KIR2DS1 were tested for binding HLA-C*02:02 or -C*05:01 pHLA multimers with and without KIR-blocking. HLA-C multimers of both subtypes showed strong binding to these cell lines and co-stained with respective KIRs, confirming the C2-group specificity of these HLA-C complexes. The HLA-C multimer binding was blocked when KIR-blocking antibody was applied before the multimer staining (Fig. 2D).

In summary, we demonstrate that KIR-blocking is an effective strategy to differentiate HLA-C multimer binding to KIR versus TCR on CD8^+^ T-cells.

### The KIR-blocking approach is dependent on HLA-C-KIR specificity

HLA-C allotypes are grouped as C1 and C2 based on a dimorphism at position 80 in the α1 domain, which determines their binding specificity to KIRs. C1 allotypes (Asn80; e.g., HLA-C*07:01 and - C*07:02) primarily interact with KIR2DL2/3, whereas C2 allotypes (Lys80; e.g., HLA-C*02:02 and -C*06:02) preferentially bind KIR2DL1/DS1/DS3/DS5 (18). As CD8^+^ T-cells can express both C1-and C2-reactive KIRs (22), C1- versus C2-reactive KIR frequencies were quantified to determine their impact on pHLA multimer-based detection of antigen-specific T-cells. Across 21 healthy donors, 4.4% (SD±3.4%) of CD8^+^ T-cells were KIR2DL2/3^+^, 1.7% (SD±3.4%) were KIR2DL1/DS1/DS3/DS5^+^, and 0.6% (SD±1.3%) co-expressed both receptor groups (Fig. 3A, B). Consistent with prior reports, these data demonstrate a higher abundance of KIR2DL2/3 compared with KIR2DL1/DS1/DS3/DS5 on CD8^+^ T-cells (27). CD8^+^ T-cells expressing either C1- or C2-reactive KIRs in the same donor showed co-staining with respective pHLA-C1 multimers (pHLA-C*07:02 multimers with KIR2DL2/L3^+^ CD8^+^ T-cells, and pHLA-C*02:02 with KIR2DL1/DS1/DS3/DS5^+^ cells), consistent with KIR-mediated binding (Fig. 3C). A small fraction of pHLA-C1 multimers showed co-staining with C2-type KIRs and vice-versa, likely reflecting binding to the CD8^+^ T-cell expressing both C1- and C2-type KIRs (Supplementary Fig. 8). In the KIR-blocking setup, pHLA-C07:02 multimer binding was specifically blocked by anti-KIR2DL2/L3 antibodies (Fig. 3D, left; Supplementary Fig. 9), whereas the binding of pHLA-C*02:02 multimer was blocked only by anti-KIR2DL1/DS1/DS3/DS5 (Fig. 3D, right). Combining both antibodies abrogated the binding of both pHLA multimer types (Fig. 3E). Thus, KIR-mediated pHLA-C multimer binding is dictated by the HLA-C-KIR specificity, requiring KIR-blocking antibodies specific to the KIRs expressed on CD8^+^ T-cells.

**Figure 3:**
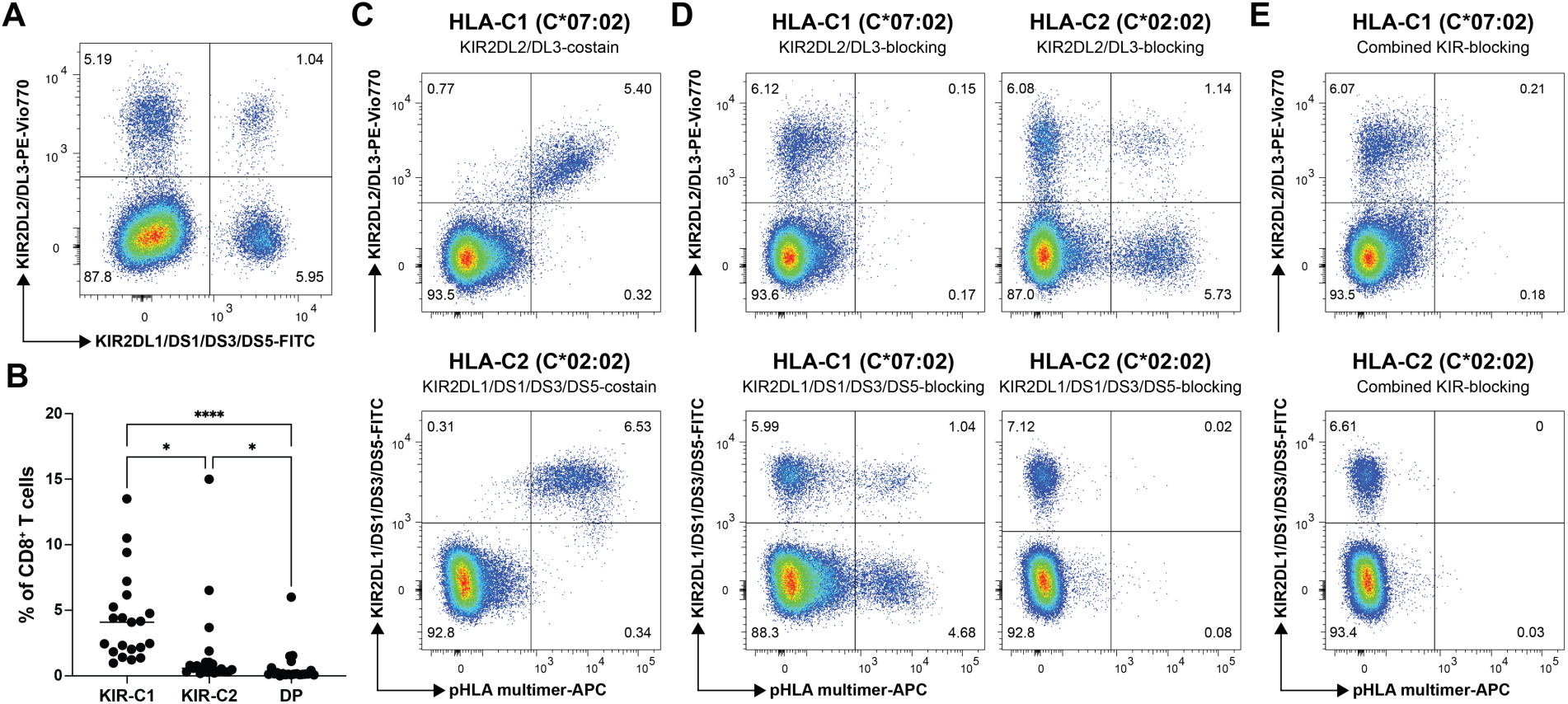
KIR-mediated pHLA-C multimer binding to CD8^+^ T-cells follows HLA-C group-specific KIR recognition. **A)** Representative flow cytometry plot showing expression of KIR2DL1/DS1/DS3/DS5 and KIR2DL2/DL3 on CD8^+^ T-cells. **B)** Frequencies of KIR-C1 (KIR2DL2/DL3), KIR-C2 (KIR2DL1/DS1/DS3/DS5), and double-positive (DP) CD8+ T-cells in healthy donors (n = 21). Groups were compared using a Friedman test followed by Dunn’s multiple-comparison test; adjusted p-values were p = 0.0164 for KIR-C1 vs. KIR-C2, p < 0.0001 for KIR-C1 vs. DP, and p = 0.0164 for KIR-C2 vs. DP. **C)** Association between HLA-C1 (C*07:02) or HLA-C2 (C*02:02) group of pHLA multimer staining and expression of the corresponding KIR group on CD8^+^ T-cells. **D)** HLA-C1 and HLA-C2 multimer binding after blockade with KIR antibodies matched or mismatched to the HLA-C group. **E)** Effect of combined KIR-C1 and KIR-C2 blockade on HLA-C multimer binding. Data in **A** and **C-E** are from the same representative donor.

### KIR- and TCR- driven HLA-C multimer binding marks distinct T-cell subsets

To determine whether CD8^+^ T-cells engaging HLA-C multimers through a TCR versus KIR represent distinct T-cell populations, we performed multimodal single-cell profiling of sorted multimer-binding cells from a healthy donor using single-cell compatible DNA-barcoded pHLA multimers (C*02:02-IACPIVMRY) under two staining conditions: multimer staining either before or after KIR antibody staining (+/– KIR-blocking) (Fig. 4A). We selected a donor in whom HLA-C*02:02–IACPIVMRY multimer binding reflected both cognate TCR-mediated recognition and KIR-mediated binding by KIR2DL1/DS1/DS3/DS5-expressing CD8⁺ T-cells, enabling us to resolve these distinct populations by single-cell analysis. The workflow combined 10x Genomics 5′ single-cell RNA sequencing, paired TCR sequencing, DNA-barcoded pHLA specificity assignment, CITE-seq using TotalSeq-C antibodies, and sample multiplexing via hashtag oligonucleotides (29). In the condition where multimer staining preceded KIR antibody staining, the sorted population included both conventional TCR-mediated multimer-binding cells and cells binding the multimer through KIR. In contrast, pre-staining with KIR antibodies prevented this interaction and thereby excluded the KIR-mediated multimer-binding population.

**Figure 4:**
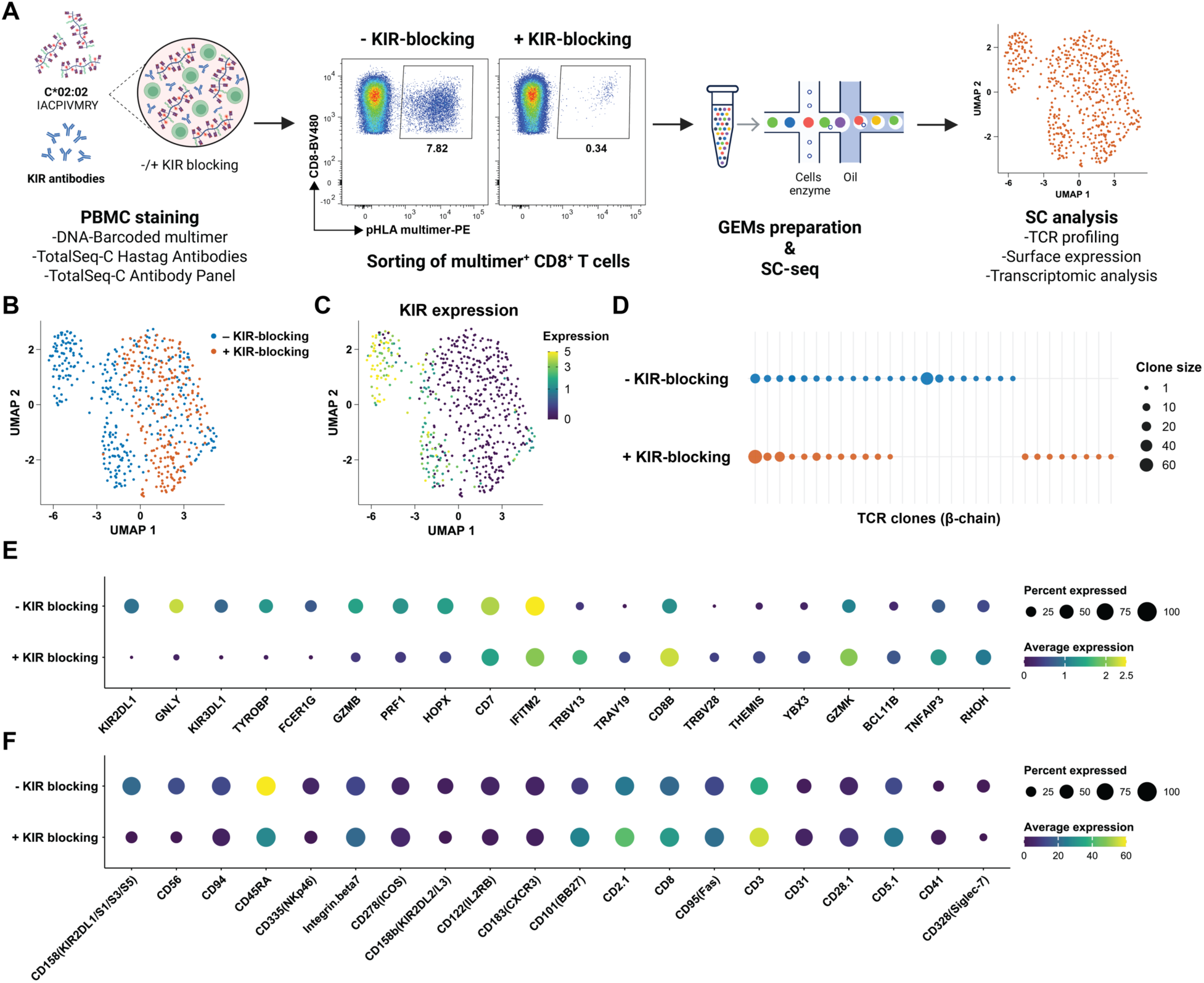
Single-cell profiling reveals distinct CD8^+^ T-cell states defined by TCR- and KIR-mediated pHLA-C multimer binding. **A)** Schematic of multimodal single-cell profiling of C*02:02-IACPIVMRY pHLA-C multimer-binding CD8^+^ T-cells sorted with or without KIR-blocking. Flow cytometry plots showing frequency of CD8^+^ T-cells binding to pHLA multimers in both the conditions. **B)** UMAP showing transcriptional clustering of sorted CD8^+^ T-cells by KIR-blocking condition. **C)** Total KIR gene expression (*KIR2DL1*, *KIR2DL2*, *KIR2DL4*, *KIR3DL1*, *KIR3DL2*, and *KIR3DL3*) projected onto the UMAP. **D)** Distribution and size of TCR clones (β-chain) of the CD8^+^ T-cells binding to pHLA multimers without and with KIR-blocking. TCR were assigned to pHLA multimers based on the pHLA-specific DNA-barcodes. Size of the dots indicate clonotype frequency based on the read counts. **E)** DEG analysis showing top 10 significantly differentially expressed genes between cells sorted with or without KIR-blocking. **F)** Similar to E, top 10 significantly differentially expressed surface proteins between cells sorted with or without KIR-blocking.

Unsupervised transcriptomic analysis revealed that T-cells from the two staining conditions segregate into partly overlapping but clearly distinct clusters on UMAP, indicating substantial compositional differences between the sorted populations (Fig. 4B). This separation corresponded closely to KIR expression, with cells from the non-blocking condition enriched in a distinct KIR-expressing cluster that was largely absent when KIR binding was blocked (Fig. 4C). Fig. 4C shows pooled KIR expression for KIR2DL1/3/4 and KIR3DL1/2/3, whereas expression data for individual KIR genes is provided in Supplementary Fig. 10. TCR clonotype analysis further supported the presence of both shared and condition-specific populations (Fig. 4D). Clonotypes detected in both with and without KIR-blocking conditions were interpreted as TCR-mediated pHLA binders, whereas clonotypes detected only in the absence of KIR-blocking were consistent with KIR-dependent multimer binding. Accordingly, shared clonotypes were excluded from the KIR-blocking population before downstream transcriptomic and protein expression analyses, enabling comparison of TCR-specific cells with CD8⁺ T cells binding pHLA-C multimers through KIRs (Fig. 4E, F). This refined analysis revealed that clonotypes lost upon KIR blockade exhibited increased KIR expression and an antigen-experienced CD8 T-cell phenotype compared with clonotypes shared across conditions (Supplementary Fig. 11). Additional unique clonotypes were also detected only in the KIR-blocking sample, likely reflecting increased sensitivity in capturing pHLA-C-TCR interactions in condition where KIR binding is abolished. Together, these data indicated that TCR-and KIR-driven HLA-C multimer interactions identify transcriptionally and clonotypically distinct CD8 T-cell populations. At the transcript level, cells without KIR-blocking showed higher expression of *HOPX, PRF1, IFITM2, CD7, GNLY*, and *TYROBP*, consistent with a cytotoxic, NK-like program (30), whereas cells from the KIR-blocking condition were enriched for TCR-associated transcripts including *TRBV13* and *TRAV19*, as well as *TNFAIP3* and *GZMK* (Fig. 4E). At the protein level, the non-blocked population was characterized by higher expression of CD45RA, CD158a, CD158b, CD56, CD335 and CD94, with CD45RA showing the strongest association, while the KIR-blocking population showed relatively higher expression of CD8, CD2.1, and CD3 (Fig. 4F). Together, these data indicated that TCR- and KIR-driven HLA-C multimer interactions identify transcriptionally, phenotypically, and clonally distinct CD8^+^ T-cell populations.

### Application of KIR-blocking strategy in large-scale detection of antigen-specific T-cells

To distinguish epitope-specific CD8^+^ T-cell binding to pHLA-C multimers from KIR-mediated binding, we integrated KIR-blocking into the DNA-barcoded pHLA multimer platform for high-throughput T-cell profiling. We applied the strategy for both HLA-C1 (HLA-C*07:01) and HLA-C2 pHLA multimers using anti-KIR2DL2/3 and anti-KIR2DL1/DS1/DS3/DS5 antibody respectively (Fig. 5A). A total of 573 unique SARS-CoV-2- and CMV-derived peptides predicted to bind HLA-C*06:02 (SARS-CoV-2, n=280; CMV, n= 7), HLA-C*07:01 (SARS-CoV-2, n=259; CMV, n=39) and HLA-C*07:02 (SARS-CoV-2, n=353; CMV, n=35) were analyzed in HLA-matched PBMCs with and without KIR-blocking (Supplementary Table 4). Screening of HLA-C*06:02-restricted peptides was performed in healthy donors (HD; n=3, collected in 2021-2022), while HLA-C*07:01- and HLA-C*07:02-restricted peptides were screened in donors with confirmed SARS-CoV-2 infection (SC2; n=8). PE- and APC-labelled multimers were used to distinguish SARS-CoV-2- and CMV-binding CD8^+^ T-cell populations respectively.

**Figure 5:**
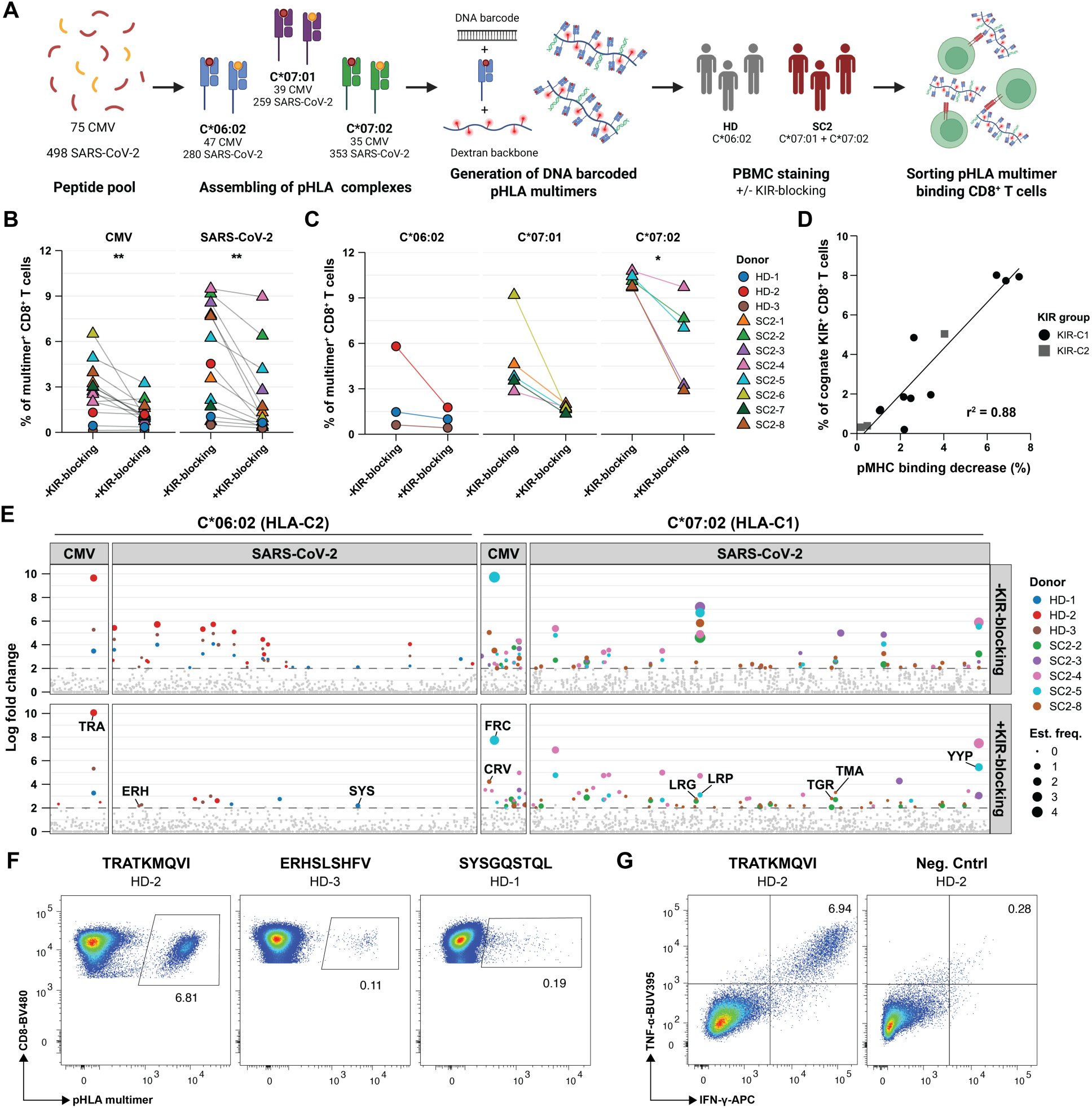
KIR-blocking enables large-scale detection of HLA-C-restricted antigen-specific CD8^+^ T-cells. **A)** Schematic overview of the DNA-barcoded pHLA multimer analysis pipeline in healthy donors (HD; n=3) and SARS-CoV-2-infected donors (SC2; n=8). HLA-C*06:02 multimers (SARS-CoV-2, n=280; CMV, n=47) were screened in HD, whereas HLA-C*07:01(SARS-CoV-2, n=259; CMV, n=39) and HLA-C*07:02 (SARS-CoV-2, n=353; CMV, n=35) multimers were screened in SC2 donors. PBMCs were incubated with DNA-barcoded pHLA multimers with or without KIR-blocking antibodies before sorting of multimer^+^ CD8^+^ T-cells. **B)** Frequency of pHLA-C multimer binding CD8^+^ T-cells across CMV and SARS-CoV-2 pools with and without KIR-blocking (Sort plots are shown in Supplementary Fig. 12 and 13). Paired donor-level comparisons are shown for each peptide pool (paired t-test; CMV, p=0.0043; SARS-CoV-2, p=0.0016). **C)** Change in pHLA multimer binding frequencies of CD8+ T-cells after KIR-blockade across HLA-C*06:02, HLA-C*07:01, and HLA-C*07:02. Paired comparisons are shown for each donor (paired t-test, p=0.0227). **D)** Association between cognate KIR expression and reduction in pHLA-C multimer binding after KIR-blocking. Each dot represents one donor-HLA combination, with colors indicating cognate KIR specificity. Linear regression analysis showed an association (r2=0.88, p<0.001). **E)** Summary of SARS-CoV-2- and CMV-derived peptides binding to CD8^+^ T-cells based on DNA-barcoded pHLA-C multimers (HLA-C*06:02 and HLA-C*07:02) with and without KIR-blocking. pHLA multimers binding to CD8 T-cells pHLA-C binders defined detection of HLA-C*06:02 and HLA-C*07:02 epitopes with and without KIR-blocking. HLA-C*06:02 represents HLA-C2, whereas HLA-C*07:02 represents HLA-C1. Significant responses were identified based on the enrichment of DNA barcodes associated with each of the tested pHLA specificities (Log2 fold change >2 and p<0.001, analyzed using Barracoda). Colored markers indicate significant donor-specific responses, with marker size reflecting estimated CD8^+^ T-cell frequency. Grey dots indicate non-significant enrichment. Selected epitopes are annotated. **F)** Conventional fluorophore labelled pHLA tetramer analysis of the three HLA-C*06:02-restrcited peptides (two SARS-CoV-2, ERHSLSHFV and SYSGQSTQL; and one CMV, TRATKMQVI) identified using DNA-barcode-based pHLA multimer analysis after KIR-blocking (Fig. 5E). PBMCs from three donors were expanded in the presence of the respective peptides and the CD8^+^ T-cell reactivity was assessed after KIR-blocking. Numbers on the plot show the identified frequency of CD8^+^ T-cells binding to each of the pHLA multimers. **G)** Representative flow cytometry plot showing functional activation (IFN-γ and TNF-α release) of expanded CD8^+^ T-cells upon stimulation with HLA-C*06:02-restrcited TRATKMQVIT peptide.

In the absence of KIR blockade, pHLA-C multimers bound a substantial fraction of CD8^+^ T-cells across the HLA-C allotypes tested (Supplementary Figs. 12 and 13). Blocking KIRs significantly reduced binding of both CMV- and SARS-CoV-2–derived pHLA-C multimers across all three HLA-C specificities analyzed, indicating that a large proportion of the observed multimer binding was KIR-mediated rather than TCR-dependent (CMV, p = 0.0043; SARS-CoV-2, p = 0.0016; Fig. 5B, 5C) This pattern was consistent with cognate KIR expression, with pHLA-C multimer binding preferentially enriched among CD8^+^ T-cells expressing the corresponding KIRs (Supplementary Fig. 14). Accordingly, the decrease in pHLA-C multimer binding after KIR-blocking strongly correlated with the frequency of cognate KIR^+^ CD8^+^ T-cells across HLA-C*06:02, HLA-C*07:01 and HLA-C*07:02 donors (r²=0.88, p<0.001; Fig. 5D).

To distinguish TCR-mediated from KIR-mediated pHLA-C multimer binding, we resolved the peptide specificity of multimer-bound CD8^+^ T-cells in the presence or absence of KIR-blocking. Without KIR blockade, several SARS-CoV-2– and CMV-derived peptide–HLA-C complexes bound CD8^+^ T-cells across all tested donors, but most such complexes either didn’t bind or showed markedly reduced binding after KIR blockade. By contrast, several specificities were retained, including known CMV epitopes restricted by HLA-C*06:02 (TRA; TRATKMQVIT) and HLA-C07:02 (FRC; FRCPRRFCF and CRC; CRCLCCYVL), indicating TCR-dependent recognitions (Fig. 5E, Supplementary Fig. 15).

Importantly, within the large SARS-CoV-2 peptide library analyzed across the three HLA-C alleles, several pHLA-C complexes showed binding in all tested donors in the absence of KIR blockade, although this pattern was not observed for all complexes. Upon KIR-blocking, however, the majority of these responses became undetectable, indicating that the observed binding was mediated predominantly by KIRs rather than by TCRs. Notably, only a subset of the tested SARS-CoV-2 peptides accounted for this KIR-mediated binding, suggesting a peptide-specific preference and hierarchy in KIR-dependent recognition of pHLA-C complexes by CD8 T-cells (Fig. 5E, Supplementary Fig. 15, Supplementary Tables 5 and 6). Together, these findings reveal that KIR-mediated pHLA-C multimer binding is not random, but highly peptide-selective, and can therefore be mistaken for TCR-mediated antigen recognition during HLA-C-restricted T-cell screening.

To further validate these findings, we selected peptides that retained pHLA-C multimer binding after KIR blockade across the three HLA-C restrictions and expanded donor-specific CD8^+^ T-cells for two weeks with individual peptides. TCR specificity was then assessed by pHLA multimer staining and cytokine release assays measuring IFN-γ and TNF-α after peptide restimulation. CD8^+^ T-cells expanded with the immunodominant HLA-C*06:02–restricted CMV epitope TRA (TRATKMQVIT; n=3), as well as two SARS-CoV-2 epitopes (ERHSLSHFV and SYSGQSTQL), showed both pHLA-C multimer binding and cytokine production, confirming TCR-specific recognition of these peptides. Similarly, three HLA-C*07:01–restricted peptides and seven HLA-C*07:02–restricted peptides induced functional activation of CD8^+^ T-cells upon peptide stimulation (Fig. 5F, 5G; Supplementary Fig. 16).

Altogether, these data show that KIR-blocking substantially reduces non-TCR-mediated pHLA-C multimer binding and reveals underlying HLA-C–restricted TCR-specific CD8^+^ T-cell responses. These findings demonstrate that KIR-mediated binding can dominate pHLA-C multimer staining and mask true TCR-specific responses unless KIR blockade is incorporated.

## Discussion

This study establishes KIR expression on CD8^+^ T-cells as a major confounding factor in HLA-C–restricted antigen discovery, demonstrating that pHLA-C multimer binding can reflect KIR interaction rather than TCR-dependent recognition. HLA-C has historically received less attention than HLA-A and HLA-B in CD8^+^ T-cell recognition, despite its documented relevance across infection, autoimmunity and cancer (6,8–13). Across viral antigens and predicted cancer-specific neoantigens, we show that pHLA-C multimers can bind a substantial fraction of CD8^+^ T-cells in a TCR-independent manner through KIR engagement, complicating the accurate identification of TCR-dependent HLA-C–restricted responses. Consistent with previous reports (22,31), the broad variation in KIR expression between donors may further influence the extent of KIR-mediated pHLA-C multimer binding.

To address this, we developed a KIR-blocking strategy that separates TCR-mediated from KIR-mediated pHLA-C binding. While post-staining detection of KIR has previously been used to identify KIR^+^multimer^+^ T-cells (23), this approach reports KIR-mediated binding rather than preventing it. In contrast, introducing KIR-blocking antibodies before pHLA staining prevented KIR–HLA-C interactions while preserving TCR-dependent detection and functional activation, including in KIR^+^ antigen-specific CD8^+^ T-cells. By combining KIR2DL1/DS1/DS3/DS5 and KIR2DL2/3 blockade, this strategy was extended across both HLA-C1 and HLA-C2 ligand groups (18,19), supporting broad implementation in HLA-C–restricted T-cell assays.

Peptide selectivity further complicates the interpretation of KIR-mediated pHLA binding. Previous work showing that approximately 40% of HLA-compatible peptides can permit HLA-C–specific KIR binding indicates that KIR recognition is less peptide-sequence specific than TCR-mediated recognition (17). Nevertheless, peptide sequence remains important, particularly at residues near the C-terminal region where P7 and P8 can modulate KIR engagement. KIR2DL2/3–HLA-C1 interactions appear more peptide-dependent than KIR2DL1–HLA-C2 interactions, further highlighting that peptide sequence can shape KIR-mediated pHLA-C binding (16,21,32). Although KIR–HLA-peptide prediction tools may help prioritize complexes with increased likelihood of KIR binding (33), experimental KIR-blocking remains necessary to distinguish KIR-mediated from TCR-mediated multimer binding in CD8^+^ T-cell assays.

Our single-cell transcriptomic, phenotypic and clonotypic analyses demonstrate that TCR- and KIR-driven pHLA-C multimer binding identify distinct CD8^+^ T-cell populations. KIR-driven binding identified cells enriched for cytotoxic and NK-like features, including FCGR3A, TYROBP, CD16 and CD56, consistent with previous reports (14,15). These data support KIR^+^CD8^+^ T-cells as a biologically distinct subset rather than only a source of staining interference. Previous work has associated these cells with regulatory functions in autoimmunity, infection and cancer, where they may limit pathogenic immune responses but also impair anti-tumor immunity through cytotoxic suppression of disease-relevant T-cells (14,15). However, their antigen specificity, suppressive mechanisms and functional heterogeneity remain incompletely defined. By separating TCR-mediated from KIR-mediated pHLA-C binding, our approach enables antigen specificity to be assigned within KIR^+^CD8^+^ T-cell populations without confounding from KIR-mediated multimer binding.

Although our data primarily address HLA-C, similar considerations may apply to HLA-A and HLA-B allotypes interacting with KIRs, including KIR3DL1 recognition of Bw4-bearing HLA molecules (34) and KIR3DL2 interactions with HLA-A3/A11 (35,36). While these interactions are often highly peptide-specific (36,37), they may still affect interpretation in high-throughput multimer screens or samples enriched for KIR^+^CD8^+^ T-cells. Together, our findings establish KIR blocking as an experimental implementation to distinguish TCR-mediated recognition from KIR-driven pHLA binding, improving the reliability of HLA-C–restricted antigen discovery while providing a framework to study antigen-specific KIR^+^CD8^+^ T-cells.

## Materials and methods

### Human samples and HLA genotyping

Peripheral blood mononuclear cells (PBMCs) from healthy blood donors were isolated from blood samples obtained through the central blood bank at Rigshospitalet, Copenhagen. PBMCs from one patient with high-risk MDS treated at the Department of Hematology and Oncology, University Hospital Mannheim, Germany, were also included. Samples were collected after written informed consent, in accordance with the Declaration of Helsinki and with approval from the relevant regional ethics committees. PBMCs were isolated by density gradient centrifugation using Leucosep tubes (Greiner Bio-One, Cat# 227288) and Lymphoprep medium (StemCell Technologies, Cat# 07861), and cryopreserved in FCS (Gibco, Cat# 10500064) containing 10% DMSO. All samples were HLA-typed by next-generation sequencing at DKMS Life Science Lab GmbH, Germany.

### Cell lines

Cell lines T1, T2, BT-549 and MDA-MB-231 were kindly provided by the Center for Cancer Immune Therapy, Copenhagen University Hospital, Denmark. The EFM192A (DSMZ, Cat#ACC258), HEK-293T (American Type Culture Collection (ATCC), Cat#CRL-11268™), and SUP-T1 (ATCC, Cat#CRL-1942™) were purchased.

### HLA expression

HLA expression was assessed at the RNA, surface protein, and immunopeptidome levels. RNA expression of HLA-A, HLA-B, and HLA-C was analyzed in 26 melanoma patients from RNA-seq data processed using TrimGalore v0.4.0 (33), Cutadapt (34), and FastQC v0.11.2 (35), quantified with Kallisto v0.23.1 (36), and reported as transcripts per million (TPM) values. Surface HLA-A, HLA-B, and HLA-C protein levels were assessed by flow cytometry in patient-derived RCC (n=4) and melanoma (n=3) tumor cell lines, with T1 and T2 cells as positive and negative controls. Cells were cultured with and without IFN-γ (250 U/mL) in R10 medium for 21 h, stained with anti-HLA-A (Creative Diagnostics, Cat# DCABH-8114), anti-HLA-B (Abcam, Cat# 212436), or anti-HLA-C (BioLegend, Cat# 400301) primary antibodies for 30 min at 4°C, followed by FITC-conjugated secondary antibody staining (Abcam, Cat# ab6785), fixation in 1% PFA (Santa Cruz Biotechnology, Cat# sc-281692), and flow cytometric analysis. Immunopeptidomes of EFM192A, MDA-MB-231, and BT-549 breast cancer cell lines were generated by W6/32-based MHC immunoaffinity chromatography (MHC-IAC) and liquid chromatography-mass spectrometry (LC-MS) as described in Viborg et al. (37). LC-MS data were processed using MaxQuant v1.5.8.3 with a peptide FDR of 1%, and peptides with rank score <2% were assigned to the highest-scoring matched HLA using NetMHCpan 4.0 (38). Peptide intensities were normalized to the mean MS1 intensity of identified MHC peptide ligands within each run. Peptides not quantified by MaxQuant were assigned the mean MS1 intensity of the 10 lowest-intensity quantified peptides from the corresponding run.

### Viral Peptide Selection

A library of 587 viral-derived peptides from SARS-CoV-2 (GenBank ID: MN908947.3), human cytomegalovirus (CMV), and Epstein-Barr virus (EBV) was used to identify HLA-C restricted responses. These 8- to 11-mer peptides were predicted to bind four HLA molecules (HLA-C*02:02, HLA-C*06:02, HLA-C*07:01, and HLA-C*07:02) using NetMHCpan 4.1 (39). Amino acid sequences of CMV proteins were obtained from the UniProt database (40) and analyzed with NetMHCpan 4.1 to generate unique 8-11 amino acid peptides covering the full protein sequences, predicting their binding affinity. A 0.5% rank threshold was used for ORF-1-derived peptides from SARS-CoV-2 and all peptides from CMV and EBV. For non-ORF-1-derived SARS-CoV-2 peptides, a 1% threshold was applied. HLA-C restricted CMV epitopes were cross-referenced with the Immune Epitope Database (IEDB) to determine whether they were novel or previously known. Published epitopes were included in the peptide libraries even if their rank exceeded 0.5%.

### Peptides

Sequences of peptides are denoted in single letter amino acid code. All peptides were purchased from Pepscan (Pepscan Presto BV, Lelystad, Netherlands) and dissolved to 10 mM in dimethyl sulfoxide (DMSO).

### HLA tissue typing, mutation analysis and neopeptide prediction

HLA typing, mutation analysis and neopeptide prediction data were derived from previously published melanoma, bladder cancer, RCC and NSCLC cohorts (24–27). Briefly, these studies used tumor and germline DNA whole-exome sequencing and tumor RNA sequencing as input for MuPeXI to identify tumor-specific mutations and nominate 9-, 10- and 11-mer peptides predicted to bind patient-specific HLA molecules (41). Neopeptide inclusion followed cohort-specific thresholds. RCC neopeptides were selected using EL%Rank<2, and melanoma and bladder cancer neopeptides using EL%Rank<0.5 and expression>0.1 TPM, all based on NetMHCpan 4.0. NSCLC neopeptides were selected using binding affinity<500 nM or EL%Rank<2 based on NetMHCpan 2.8 or NetMHCpan 4.0. If fewer than 200 neopeptides met these criteria, the top 200 highest-ranked peptides were included.

### Production of HLA class I molecules

All HLA molecules were produced by expressing HLA class I heavy chain and human β_2_-microglobulin light chain in *Escherichia coli* (*E. coli*). Refolding was performed in the presence of UV-sensitive HLA-specific peptide ligands (42,43) or empty peptide-receptive HLA class I molecules (44). Folded monomeric HLA class I molecules were biotinylated using the BirA biotin-protein ligase standard reaction kit (Avidity LLC, Aurora, USA) and purified by size-exclusion chromatography using HPLC (Waters Corporation, USA). Before storage at -80°C, monomers were quality-controlled for concentration, UV-mediated peptide degradation and biotinylation efficiency.

### Generation of fluorescently labelled pHLA tetramers

Individual HLA-restricted pHLA monomers were generated by incubating 200 μM peptide with 100 μg/mL HLA class I molecules at for 1 h, using either UV-mediated peptide exchange or direct loading in the case of empty-loadable HLA class I molecules. Fluorescent pHLA tetramers were produced by conjugating pHLA monomers to streptavidin-fluorochrome conjugates at a final concentration of 18 μg/mL for 30 min at 4°C, followed by addition of d-biotin to a final concentration of 25 μM. Streptavidin conjugates are listed in Supplementary Table 7. For dual-color tetramers, monomers were tetramerized separately with two streptavidin conjugates, and the resulting tetramers were mixed at a 1:1 ratio before addition of freezing medium. Tetramers were supplemented with 10x freezing medium, yielding final concentrations of 0.5% BSA and 5% glycerol, and stored until use.

### pHLA multimer staining

PBMCs were thawed, washed in R10 medium (RPMI 1640 with GlutaMAX, Gibco, Cat# 61870010, supplemented with 10% FCS), and washed twice in FACS buffer (PBS with 2% FCS). Cells were incubated with 1 μL pHLA tetramer and 50 nM dasatinib for 15 min at 37°C. Antibody cocktail and BD Horizon Brilliant Stain Buffer Plus were then added, and cells were stained for 30 min at 4°C in a final volume of 100 μL (BD Biosciences, Cat# 566385; Supplementary Table 8). Data were acquired on an LSRFortessa flow cytometer (BD Biosciences).

### KIR staining of PBMCs

To investigate the role of KIRs in pHLA multimer interactions, KIR antibodies were applied either before or after pHLA multimer staining. For multimer-first staining, thawed PBMCs were first stained with pHLA multimers, followed by staining with antibodies against KIR2DL1/DS1/DS3/DS5 (HP-MA4) and/or KIR2DL2/DL3 (DX27) together with additional antibodies for cell surface markers. For KIR-blocking, KIR antibodies were added before pHLA multimer staining and incubated in FACS buffer for 30 min at either 4°C or room temperature. Antibody details and titration data for fluorochrome-conjugated and purified antibodies are provided in Supplementary Table 9.

### In vitro T-cell expansion

PBMCs were cultured for 13 days in X-VIVO™ serum-free medium (Lonza, Cat#BE02-60Q) supplemented with 5% human serum (Gibco, Cat#1027-106), 200 IU/mL IL-2, 20 ng/mL IL-15, and 1 μg/mL peptide of interest.

### T-cell functional analysis

T-cell functional activation was assessed via intracellular cytokine staining for IFN-γ and TNF-α using the eBioscience™ FoxP3/Transcription Factor Staining Buffer Set (ThermoFisher, Cat#00-5523-00). Thawed or expanded PBMCs were rested for 24 h before stimulation in X-VIVO serum-free medium supplemented with 5% human serum. Cells were stimulated for 8-10 h at 37°C with peptide at 1 μg/mL in the presence of GolgiPlug (BD Biosciences, Cat#555029, 1/1000 dilution). For samples requiring KIR-blocking, KIR2DL1/DS1/DS3/DS5-purified antibody was added and incubated for 30 min at RT prior to stimulation. Negative controls were treated with DMSO at a concentration matching peptide-stimulated samples, whereas Leukocyte Activation Cocktail was used as a positive control (BD Biosciences, Cat# 550583, 1/500 dilution). After stimulation, cells were washed twice in FACS buffer and stained for surface markers for 30 min at 4°C. Cells were then fixed in FoxP3 Fixation/Permeabilization working solution at 4°C for up to 18 h and stained intracellularly for IFN-γ and TNF-α in permeabilization buffer. Samples were washed, resuspended in FACS buffer, and analyzed by a LSRFortessa flow cytometer (BD Bioscience).

### T-cell lines expressing KIR2DS1 or KIR2DL1

Customized plasmids encoding KIR2DS1 or KIR2DL1 (GenScript USA Inc.) were amplified in E. coli One Shot Stbl3 Competent (Invitrogen, Cat# 7373-03) under carbenicillin selection, purified using the QIAGEN Plasmid Plus Midi Kit (Qiagen, Cat# 12941), and quantified using a NanoDrop 1000 spectrophotometer (Thermo Scientific). Lentiviral particles were produced in HEK-293T-cells by co-transfection of the corresponding transfer vector with pRSV.REV, pMDLg/p.RRE, and pMD2.G packaging plasmids (Addgene, Cat# 12253, 12251, and 12259) using Lipofectamine 3000 (Invitrogen, Cat# L3000001) in Opti-MEM I (Gibco, Cat# 31985062). Viral supernatants were harvested after 24 h, concentrated using Lenti-X Concentrator (Takara, Cat# 631232), and stored at **-**80°C. SUP-T1 cells were seeded at 1 × 10⁶ cells/mL in R10 medium and transduced with concentrated lentiviral particles. After 48 h, cells were expanded in fresh R10 medium. On day 5 post-transduction, cells were washed twice and tested for residual lentivirus using the Lenti-X GoStix Plus Test (Takara Bio, Cat# 631280). Successful transduction was confirmed by GFP expression using flow cytometry.

### Staining of antigen-specific T-cells using DNA-barcoded pHLA multimers

DNA-barcoded pHLA multimer libraries containing neoantigens, SARS-CoV-2-, CMV- and EBV-derived epitopes were generated as described by Bentzen et al. (4). Briefly, peptide-loaded pHLA complexes were conjugated to dextran backbones carrying unique DNA barcodes. SARS-CoV-2-derived multimers were labelled with PE, whereas CMV/EBV-derived multimers were labelled with APC. PBMCs were thawed, washed twice in R10 medium and once in barcode cytometry buffer (BCB; PBS with 0.5% BSA, 100 µg/mL herring DNA and 2 mM EDTA). Where indicated, staining was performed with or without KIR-blocking as described above. Cells were incubated with HLA-matched SARS-CoV-2, CMV and EBV DNA-barcoded pHLA multimers for 30 min at room temperature, followed by phenotyping antibody staining for 30 min at 4°C (Supplementary Table 8). Cells were washed twice in BCB, fixed in 1% PFA, washed and resuspended in BCB. Multimer-binding CD8^+^ T-cells were sorted on a BD FACSAria Fusion flow cytometer (BD Biosciences; Supplementary Fig. 9) or BD FACSDiscover S8 cell sorter (BD Biosciences; Supplementary Fig. 13). Sorted cells were centrifuged at 5.000g for 10 min, and cell pellets were stored at -20°C.

### Sequencing analysis of DNA barcodes

DNA barcodes from sorted cell pellets and a 10.000-fold diluted aliquot of the multimer pool were amplified using the Taq PCR Master Mix Kit (Qiagen, Cat# 201443). PCR products were purified using the QIAquick PCR Purification Kit (Qiagen, Cat# 28104) and sequenced by PrimBio (USA). Sequencing data were analyzed using the Barracoda 2.0 software package (https://services.healthtech.dtu.dk/tuba/barracoda-2.0/) as previously described by Bentzen et al. (4). DNA barcodes with p<0.001 and Log2 fold change (LogFC)>2 relative to baseline were considered significant and interpreted as antigen-specific T-cell responses.

### T-cell staining and sorting for single-cell analysis

PBMCs were thawed, washed in Cell Staining Buffer (PBS with 0.5% BSA), and stained with PE-labeled DNA-barcoded pHLA multimers for 60 min at 4°C. Cells were subsequently incubated with Human TruStain FcX Fc Blocking reagent (BioLegend, Cat#42302) for 10 min at 4°C, followed by staining with the TotalSeq-C Human Universal Cocktail (BioLegend, Cat#399905), TotalSeq-C hashtag antibodies, and the antibody panel listed in Supplementary Table 8. Antigen-positive CD8^+^ T-cells were sorted based on PE-labeled pHLA multimer binding using a FACS Melody cell sorter (BD Biosciences). Approximately 17,000 sorted cells were pooled and processed using the Chromium Next GEM Single Cell 5ʹ Reagent Kits v2 with Feature Barcode technology for Cell Surface Protein and Immune Receptor Mapping (10x Genomics), according to the manufacturer’s protocol. Gene expression, TCR V(D)J, antibody-derived tag, hashtag, and pHLA barcode libraries were generated, quantified using the Qubit dsDNA HS Assay Kit (Invitrogen, Cat#Q32851), pooled, and sequenced by Novogene on an Illumina NovaSeq platform with 150-bp paired-end reads.

### Single-cell RNA and ADT sequencing data analysis

Raw FASTQ files were processed with Cell Ranger using the cellranger multi pipeline and the GRCh38 reference genome. Gene expression and antibody-derived tag count matrices, together with hashtag oligo, DNA-barcoded pHLA multimer and TCR V(D)J information, were imported into Seurat v5.4.0 for downstream analysis. TCR V(D)J information was processed using scRepertoire before integration with the Seurat metadata. Hashtag oligos were CLR-normalized and used for demultiplexing with HTODemux, retaining singlet T-cells. Cells expressing <200 or >2,000 genes or ≥5% mitochondrial genes were excluded. Gene expression data were normalized using LogNormalize, and ADT data were CLR-normalized.

PCA was performed in Seurat using the top 1,800 highly variable genes, and the first 20 principal components were used for clustering and UMAP visualization. KIR RNA expression was summarized from normalized expression of KIR2DL1, KIR2DL3, KIR3DL1, KIR3DL2, KIR3DL3 and KIR2DL4, while KIR protein expression was summarized from CLR-normalized ADT expression of CD158 (KIR2DL1/S1/S3/S5), CD158b (KIR2DL2/L3) and CD158e1 (KIR3DL1). Shared TRB clonotypes were removed from the non-blocked condition before differential RNA and ADT expression analysis, while all cells from the KIR-blocked condition were retained. Differential expression was assessed using FindAllMarkers, with positive markers defined by adjusted p-value <0.05 and Log2 fold change >0.25. The top 10 RNA genes and ADT markers per condition were selected by adjusted p-value after excluding ribosomal genes and isotype/control ADT markers and ordered by a combined effect-significance score. Violin plots were generated from normalized RNA expression or CLR-normalized ADT expression, and heatmaps from scaled KIR gene expression values.

### TCR Profiling Data Analysis

TCR clonotypes were defined by the TRB CDR3 sequence, and cells without a detectable TRB CDR3 sequence were excluded. Clone size was calculated within each staining condition and pHLA barcode assignment, with clonotypes represented by more than one cell retained for visualization. For clonotype-stratified KIR expression analyses, clonotypes were classified as shared, block-sensitive, or block-only based on detection in both conditions, only without KIR-blocking, or only after KIR-blocking, respectively.

### Flow cytometry analysis

Flow cytometry data were analyzed using FlowJo v10.10.0 (FlowJo LLC). KIR expression on CD8^+^ T-cells and NK cells was evaluated as shown in Supplementary Fig. 1, and IFN-γ^+^TNF-α^+^ cells were quantified as shown in Supplementary Fig. 2. For combinatorial pHLA tetramer staining, single live FITC-negative CD8^+^ lymphocytes were gated, and antigen-specific T-cells were identified by dual positivity for the corresponding tetramer-color combination and negativity for all other tetramer colors, as shown in Supplementary Fig. 6 (2). pHLA multimer-positive CD8^+^ T-cell analysis and sorting were performed as shown in Supplementary Fig. 9 and 14.

### Data processing and statistical analysis

Data visualization and statistical analyses were performed using R version 4.5.2 (45), ggplot2 version 4.0.1 (46), and GraphPad Prism version 10.6.0. Associations between variables were assessed by simple linear regression, with r^2^ and p values reported where relevant. Paired comparisons between matched samples were analyzed using two-sided Wilcoxon matched-pairs signed-rank tests or paired t-tests, as indicated in the figure legends. Comparisons between more than two paired groups were performed using Friedman tests followed by Dunn’s multiple-comparison tests. Unpaired comparisons were analyzed using two-sided Mann-Whitney tests. Proportional differences were evaluated using two-sided Fisher’s exact tests. For single-cell clonotype-stratified analyses, two-sided Wilcoxon rank-sum tests were used with Benjamini–Hochberg correction for multiple testing. Relative expression was calculated as the ratio to the negative control where indicated. Schematic illustrations were created with BioRender.

## Acknowledgments

We are grateful to all patients who participated in the study and provided the samples included in the analysis. We thank S. Sebbaha, A. G. Burkal, T. M. L. Nguyen, B. Rotbøl, and A. F. Løye for their excellent technical assistance and all collaborators for their valuable contributions to the study.

## Author contributions

A.H.K. designed and performed experiments, analyzed data, prepared figures, and wrote the manuscript. J.K. contributed to experimental design, performed experiments, analyzed data, prepared figures, and contributed to writing the manuscript. S.R. performed experiments and analyzed data. A.B. analyzed data. Y.B., A.O.M., T.R., and M.K. performed experiments. S.R.H. conceived, designed, and supervised the study. S.K.S. conceived, designed, and supervised the study, analyzed data, and wrote the manuscript.

