## Supplementary Figures for "KIR Expression Defines a Transcriptionally Distinct CD8+ T-cell Population that Confounds Antigen-Specific T-cell Detection"

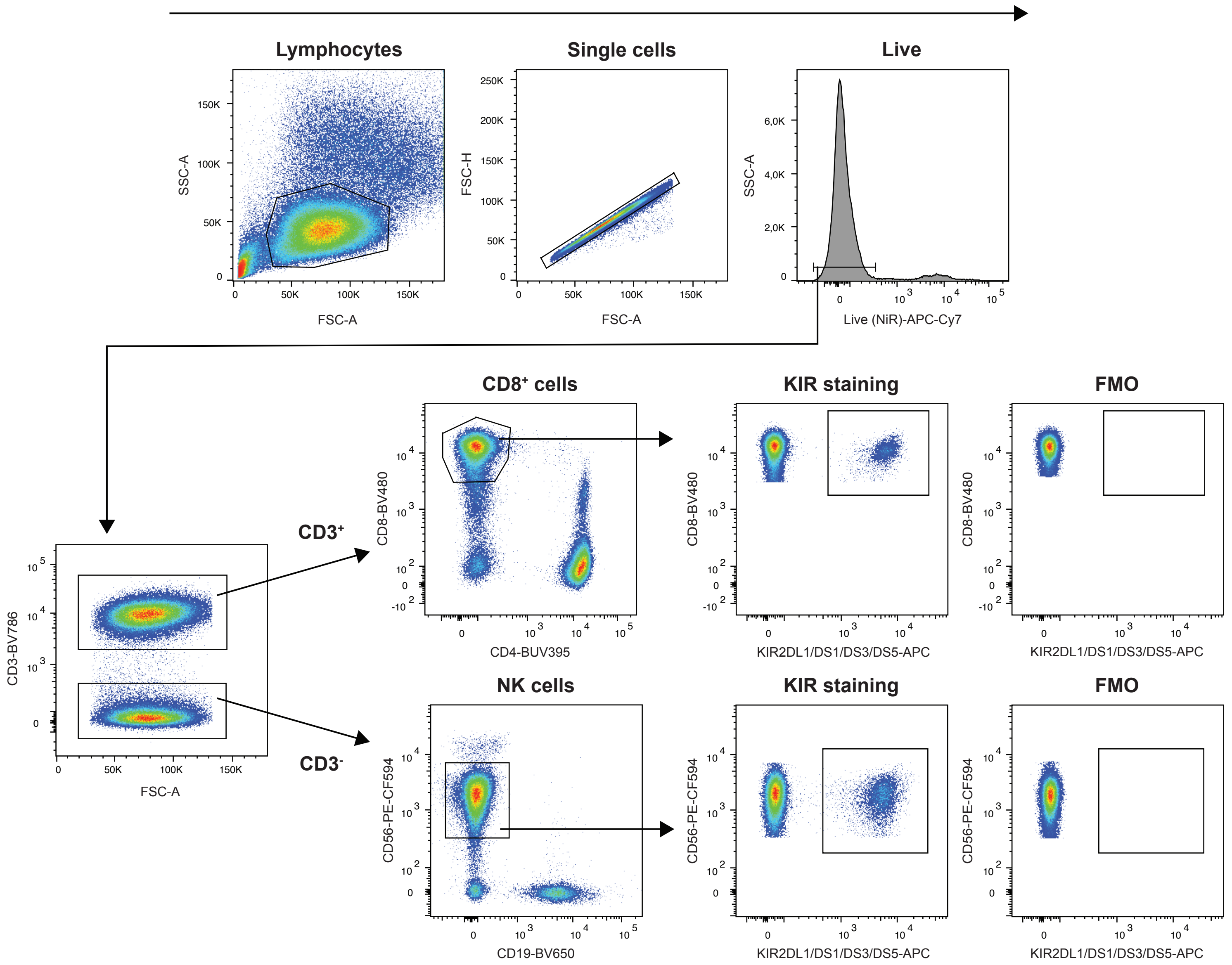


**Supplementary Figure 1: Gating strategy for KIR2DL1/DS1/DS3/DS5 expression in CD8⁺ T-cells and NK cells.**

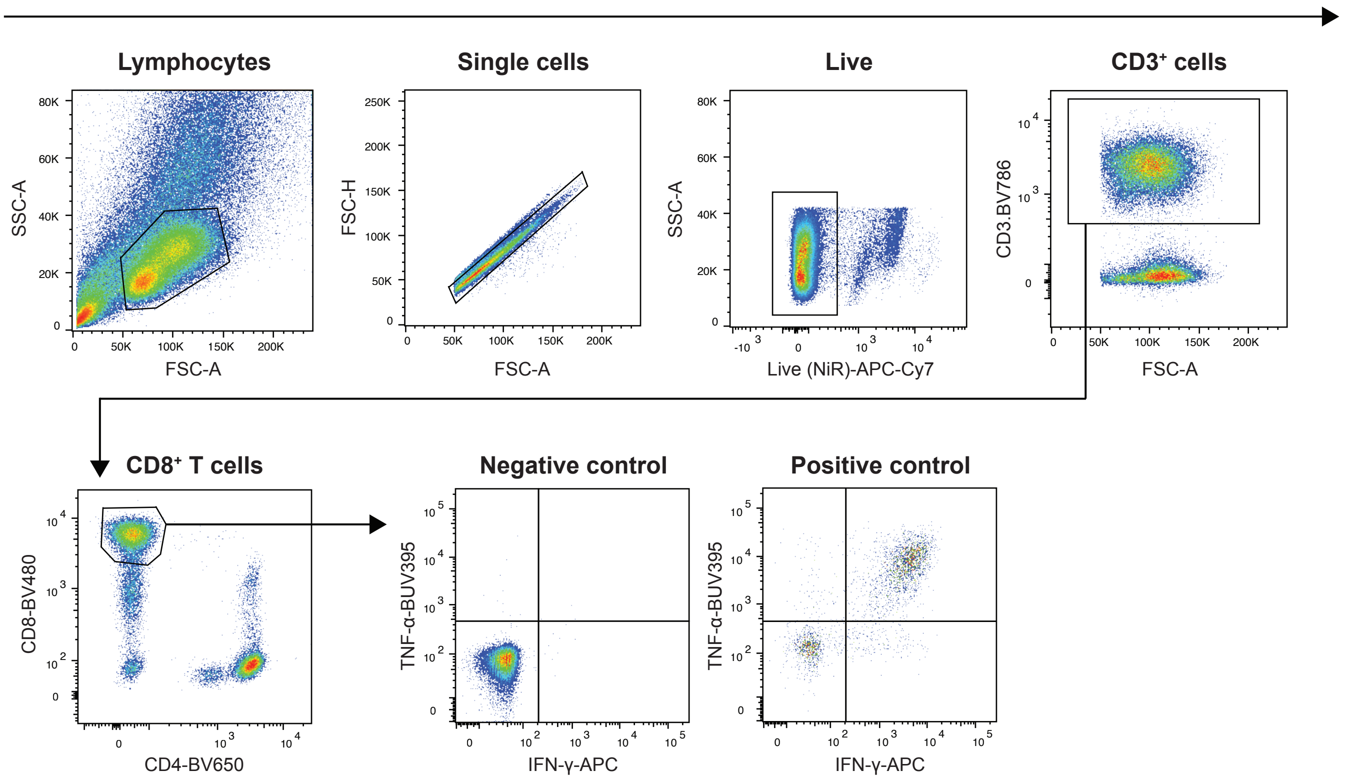


**Supplementary Figure 2: Gating strategy to assess functional activation of T-cells via cytokine release (IFN-y and TNF-a)**

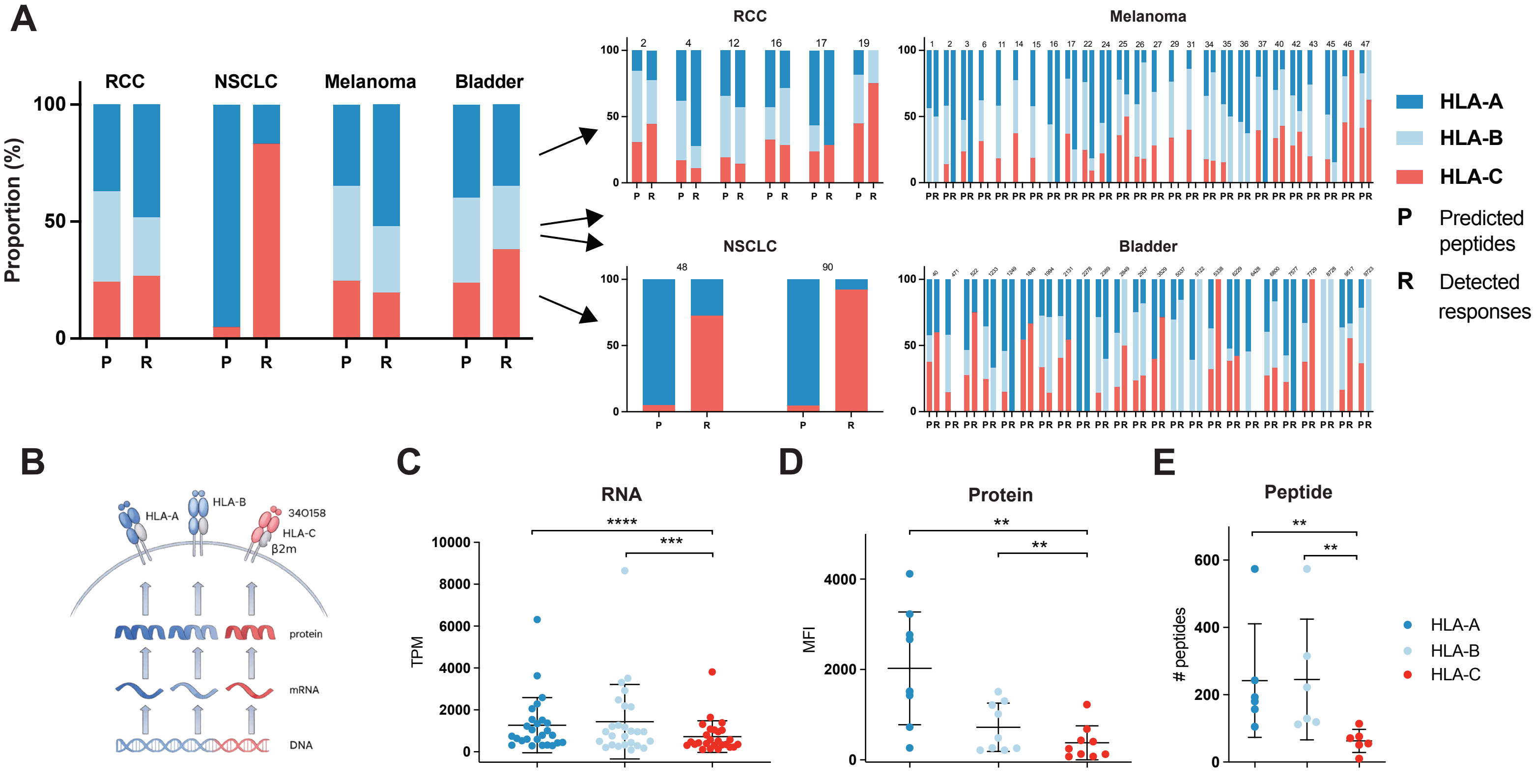


**Supplementary Figure 3: pHLA-C multimer analysis overrepresents HLA-C-restricted neoepitope responses in cancer.**
**A)** Proportional contribution of HLA-A-, HLA-B- and HLA-C-restricted neoepitopes to CD8$^{+}$ T-cell responses across four cancer cohorts. *Left*, pooled cohort analysis. *Right*, cohort-level analysis with patient-specific data comparing predicted (P) and detected (R) responses. P, predicted HLA-A/B/C neopeptides. R, detected neoepitope responses. **B)** Schematic overview of HLA-A, HLA-B and HLA-C expression assessed at RNA, protein and peptide levels. **C)** HLA-A, HLA-B and HLA-C RNA expression in malignant melanoma patients (n=26), reported as transcripts per million (TPM). Two-tailed paired Wilcoxon test (HLA-A vs. HLA-C, p<0.0001; HLA-B vs. HLA-C, p=0.0006). **D)** HLA-A, HLA-B and HLA-C surface protein expression assessed by flow cytometry in patient-derived RCC and melanoma cell lines (n=7). Each dot represents the mean of two IFN-γ-stimulated and two unstimulated samples. Two-tailed paired Wilcoxon test (HLA-A vs. HLA-C, p=0.0039; HLA-B vs. HLA-C, p=0.0039). **E)** Endogenous HLA-A-, HLA-B- and HLA-C-associated peptides measured by mass spectrometry in EFM192A, MDA-MB-231, and BT-549 breast cancer cell lines (n=3). Two-tailed unpaired Mann-Whitney test (HLA-A vs. HLA-C, p = 0.0043; HLA-B vs. HLA-C, p=0.0043).

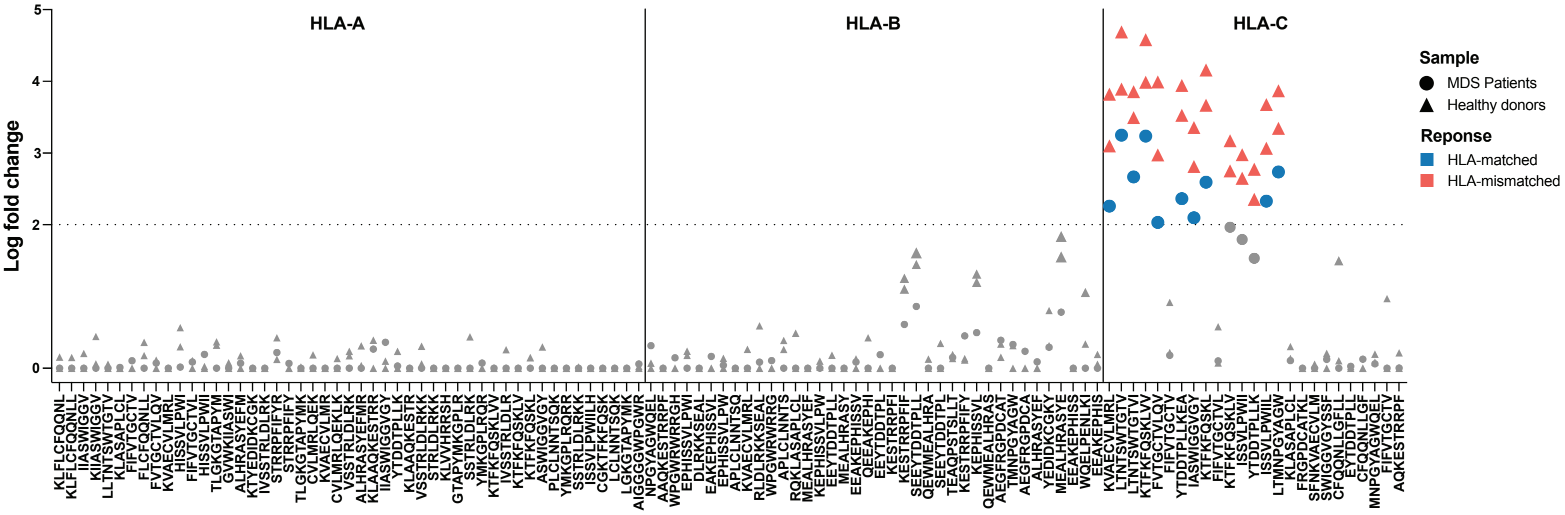


**Supplementary Figure 4: DNA-barcoded pHLA multimer screening of an MDS patient-specific neopeptide library.**
DNA-barcoded pHLA multimer screening of an MDS patient-specific neopeptide library in the index MDS patient and two healthy donors. The MDS patient carried HLA-C*02:02 and HLA-C*07:02. One healthy donor shared HLA-C*02:02 but lacked HLA-C*07:02, whereas the second healthy donor carried neither allele. DNA barcode enrichment is shown as Log2 fold change and grouped by predicted HLA restriction. Symbols indicate sample origin. Colored markers indicate significant HLA-matched or HLA-mismatched responses, whereas grey markers indicate non-significant enrichment. The dotted line marks the enrichment threshold used to define significant responses (Log2 fold change >2 and p<0.001). The full list of MDS-derived neopeptides is provided in Supplementary Table 2.

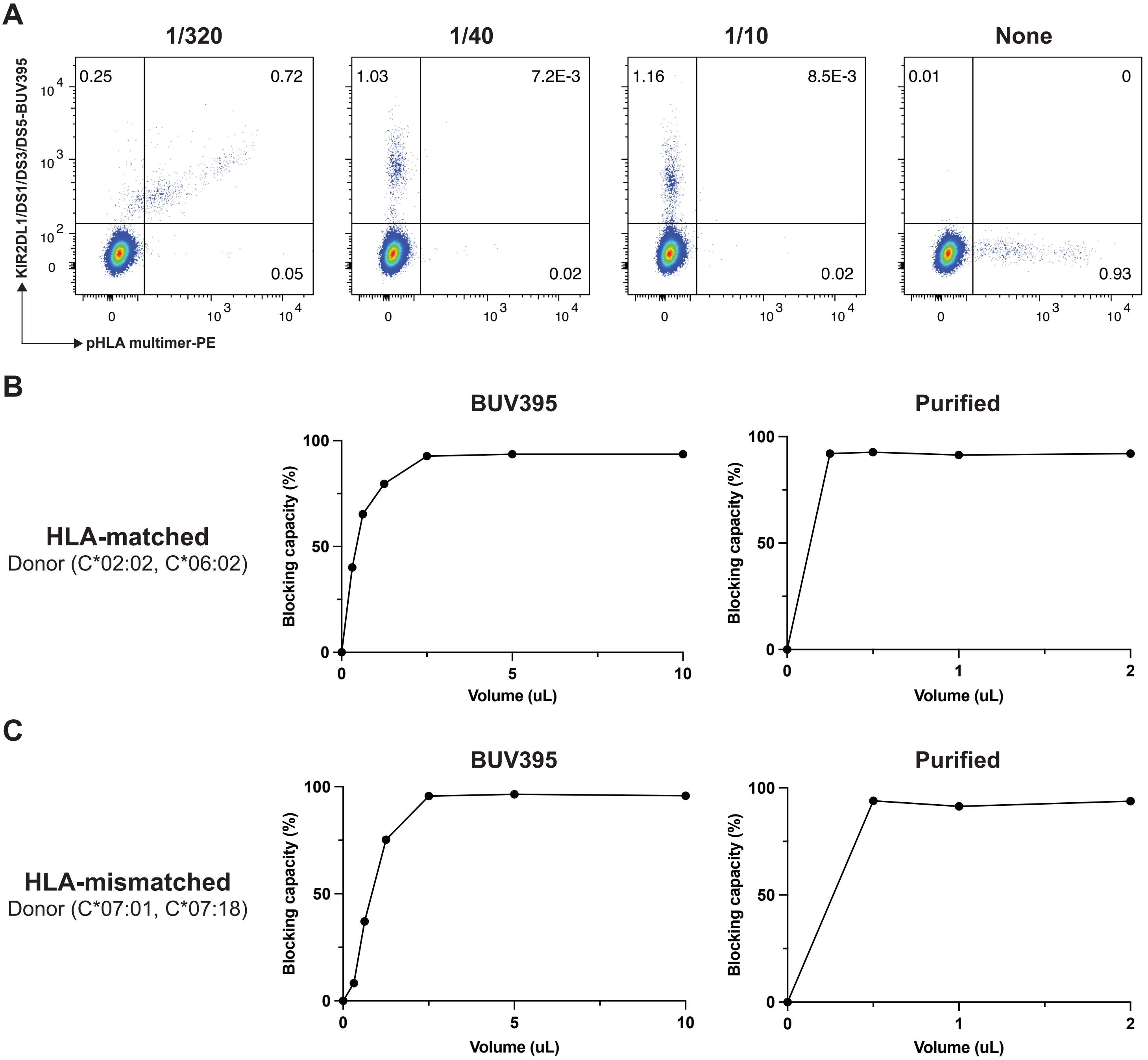


**D**

| Antibody | Recommended dilution | Manufacture |
|---|---|---|
| **KIR2DL1/DS1/DS3/DS5 (HP-MA4)** | | |
| Purified | 1/200 | BioLegend |
| BUV395 | 1/40 | BD Biosciences |
| APC | 1/40 | ThermoFisher |
| FITC | 1/40 | BioLegend |

**Supplementary Figure 5: Titration of KIR2DL1/DS1/DS3/DS5 blocking antibodies for inhibition of HLA-C*02:02-IACPIVMRY pHLA-C multimer binding to CD8⁺ T-cells.**

**A)** Flow cytometry plots showing HLA-C*02:02-IACPIVMRY multimers binding after adding anti-KIR2DL1/DS1/DS3/DS5 antibodies at different concentrations. **B)** Titration plots showing efficiency of KIR-blocking using BUV395 fluorophore conjugated antibody (*left*) and antibody without any fluorophore (*right*) in an HLA-C*02:02-matched donor. **C)** Similar to B, antibody titration results in an HLA-C*02:02-mismatched donor. *Left*, using BUV395 fluorophore conjugated antibody and *right*, antibody without any fluorophore. **D)** Summary of recommended dilutions of different fluorophore conjugated and non-fluorophore conjugated KIR2DL1/DS1/DS3/DS5 antibodies for KIR-blocking.

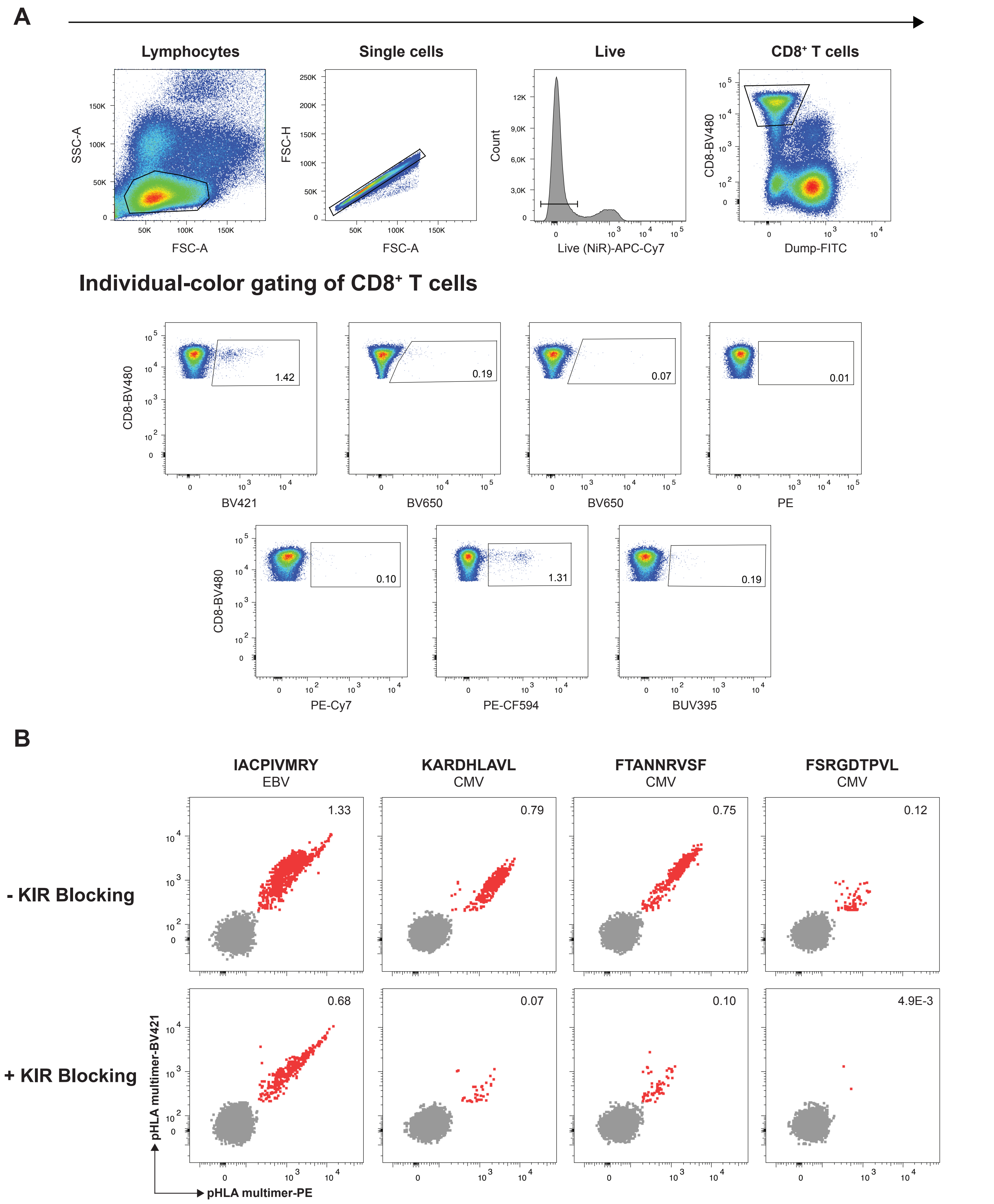


**Supplementary Figure 6: HLA-C*02:02-restricted CMV- and EBV-derived responses identified by combinatorial pHLA multimer analysis.** **A)** Flow cytometry gating strategy used to identify combinatorially encoded pHLA-C multimer⁺ CD8⁺ T-cell responses. PBMCs from two HLA-C*02:02-matched healthy donors were screened using pHLA-C multimers representing CMV- and EBV-derived peptides predicted to bind HLA-C*02:02. The 46-peptide library was divided into four screening panels containing up to 14 peptides each. Individual peptide specificities were identified by unique two-color combinations of PE-CF594-, BUV395-, BV421-, BV605-, BV650-, and PE-Cy7-labeled multimers. Representative flow cytometry plots from one donor are shown. **B)** Validation of responses identified by combinatorial encoding using individual pHLA-C tetramer staining (dual fluorophore labelled) with or without KIR-blocking (anti-KIR2DL1/DS1/DS3/DS5).

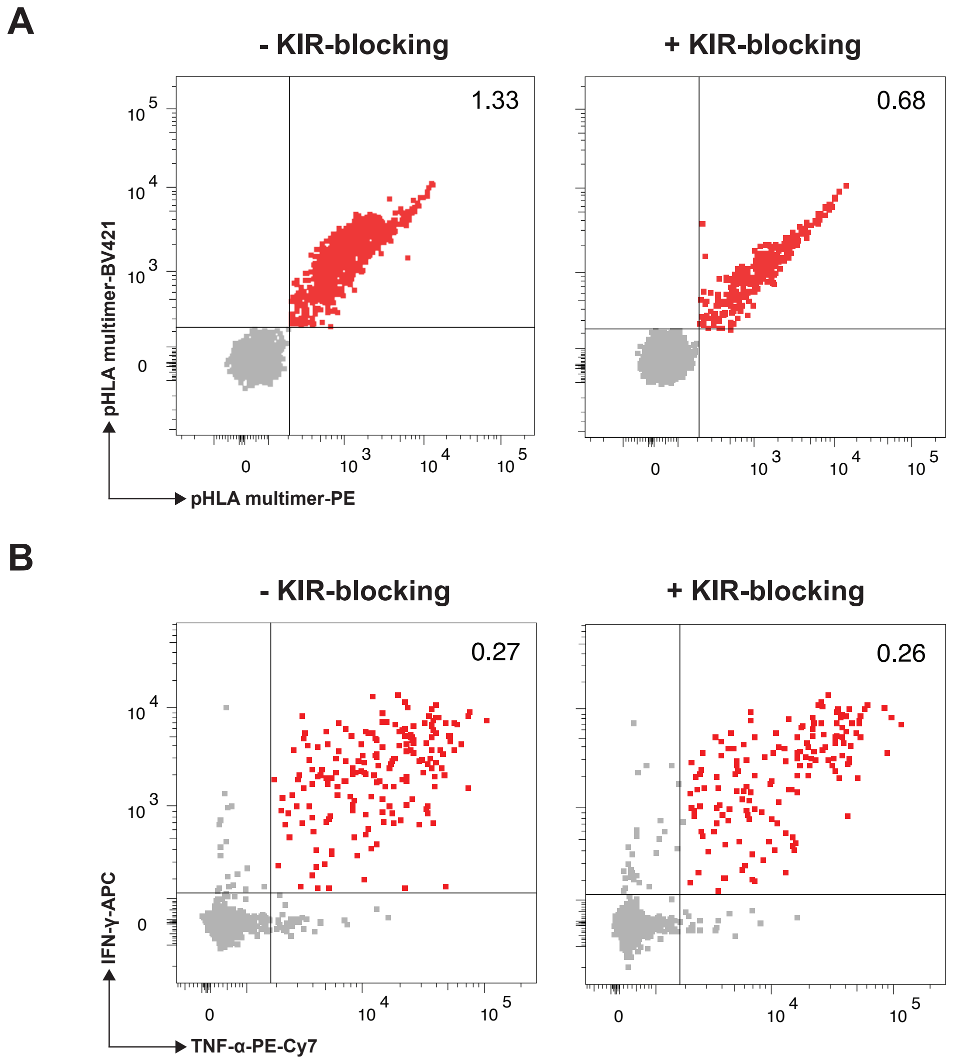


**Supplementary Figure 7: KIR-blocking separates KIR-mediated pHLA-C binding without interfering with TCR-driven activation.**
**A)** HLA-C*02:02-IACPIVMRY pHLA-C tetramer staining of $CD8^+$ T-cells with or without KIR-blocking in a donor with both TCR- and KIR-mediated binding. **B)** Functional activation of $CD8^+$ T-cells assessed by IFN-γ and TNF-α release following IACPIVMRY stimulation with or without KIR-blocking in the same donor.

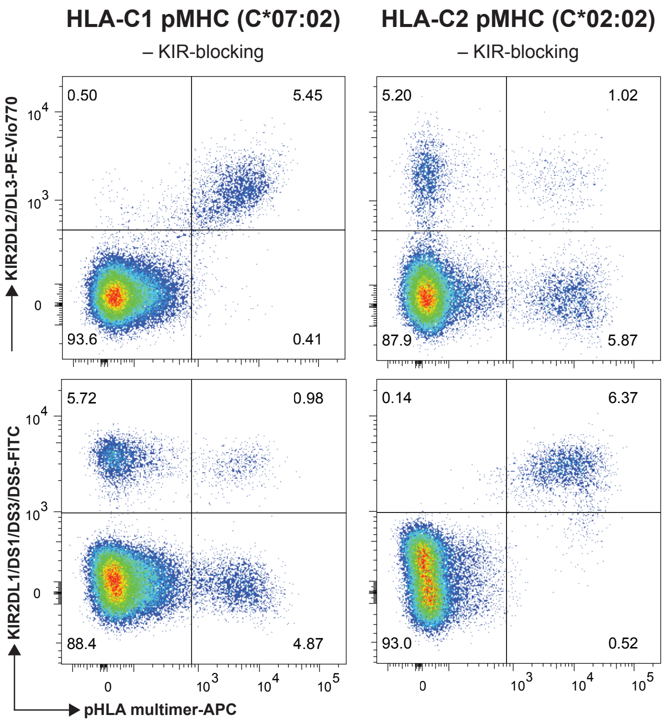


**Supplementary Figure 8: HLA-C1 and HLA-C2 pHLA multimers preferentially bind their corresponding KIR populations.**
Representative flow cytometry plots showing HLA-C*07:02 (HLA-C1) and HLA-C*02:02 (HLA-C2) pHLA multimer binding relative to KIR2DL2/DL3 and KIR2DL1/DS1/DS3/DS5 expression, respectively, in $CD8^+$ T-cells in the absence of KIR-blocking.

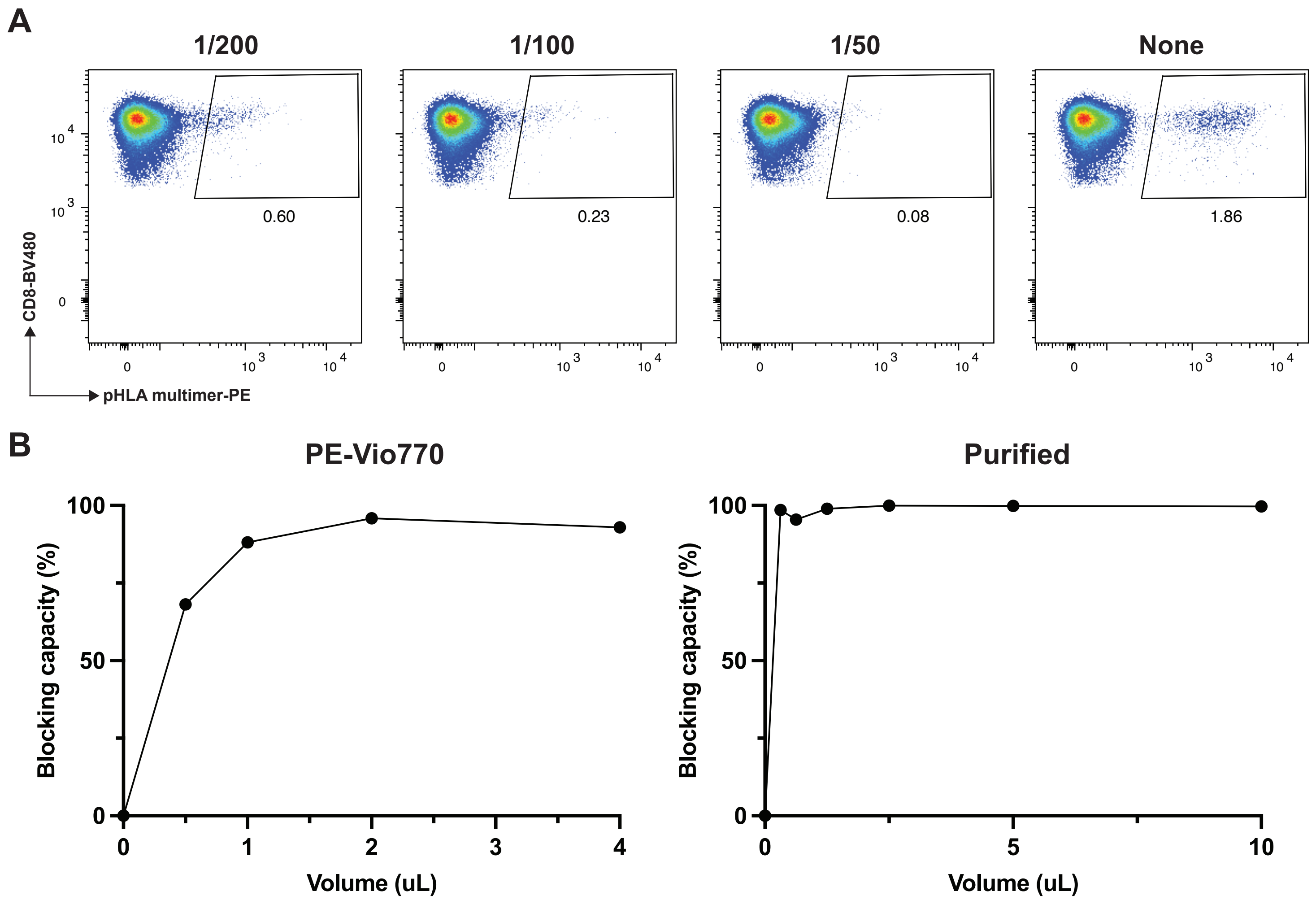


C

| Antibody | Recommended dilution | Manufacture |
|---|---|---|
| **KIR2DL2/DL3 (DX27)** | | |
| Purified | 1/50 | Miltenyi Biotec |
| PE-Vio770 | 1/50 | Miltenyi Biotec |

**Supplementary Figure 9: Titration of KIR2DL2/DL3 blocking antibodies for inhibition of HLA-C*07:02-QRNAPRITF pHLA-C tetramer binding.**

**A)** Flow cytometry plots showing HLA-C*07:02-QRNAPRITF pHLA-C tetramer binding to $CD8^+$ T-cells following blocking with different concentrations of anti-KIR2DL2/DL3 (PE-Vio770 conjugated) antibody. **B)** Titration curves of KIR-blocking using a fluorophore conjugated (*left*, PE-Vio770) and non-conjugated (*right*, Purified) anti-KIR2DL2/DL3 antibody in an HLA-C*07:02-matched donor without a detectable TCR-mediated response to QRNAPRITF. **C)** Titration results for PE-Vio770-conjugated and purified KIR2DL2/DL3 antibodies.

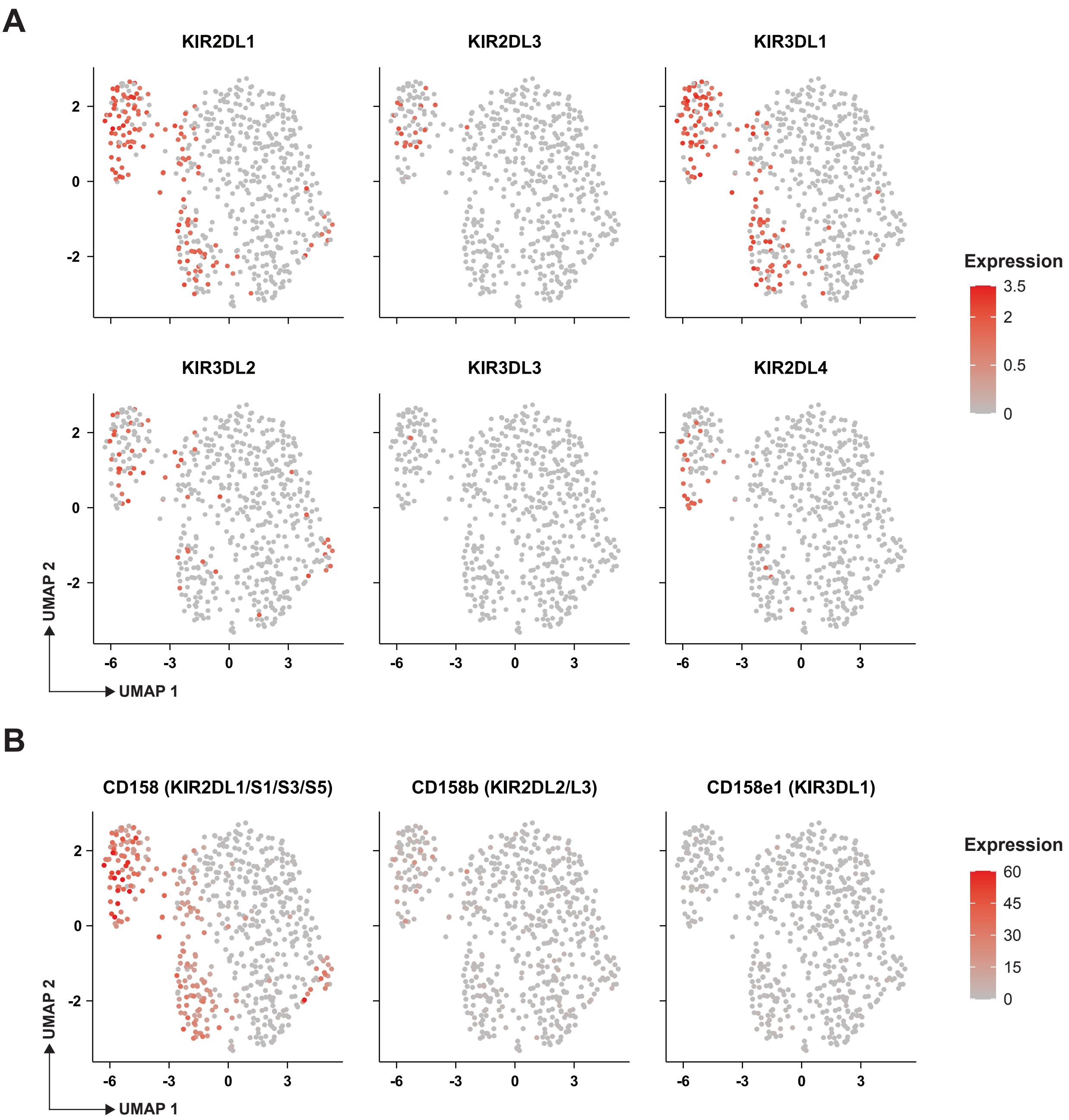


**Supplementary Figure 10: KIR gene and surface protein expression projected onto single-cell UMAPs.**
**A)** Expression of *KIR2DL1*, *KIR2DL3*, *KIR3DL1*, *KIR3DL2*, *KIR3DL3*, and *KIR2DL4* transcripts projected onto the single-cell RNA-sequencing UMAP. **B)** Surface expression of CD158 (KIR2DL1/DS1/DS3/DS5), CD158b (KIR2DL2/DL3), and CD158e1 (KIR3DL1) projected onto the CITE-seq UMAP.

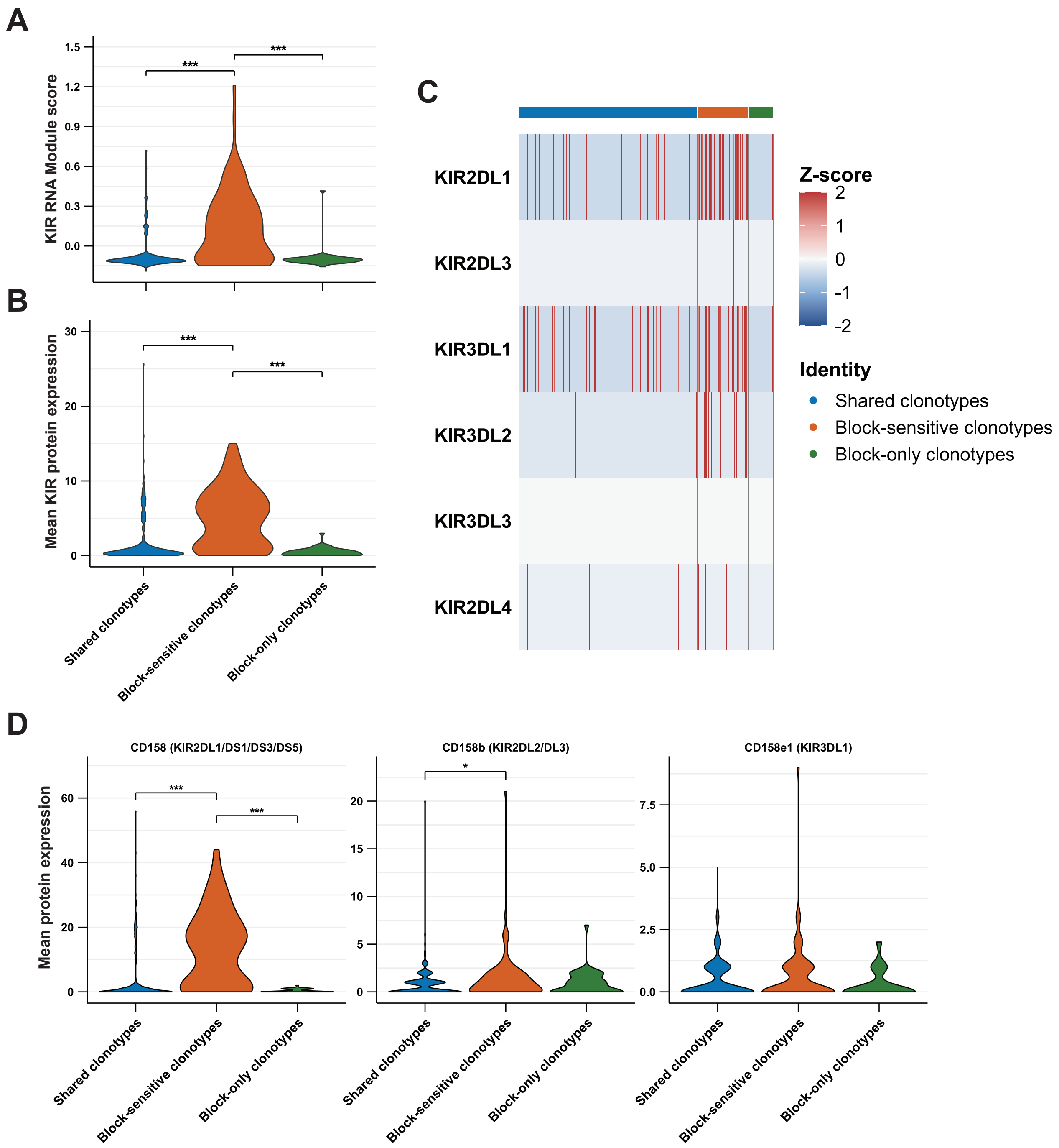


**Supplementary Figure 11: KIR expression stratified by TCR clonotypes.**
Clonotypes were classified as shared, detected both without and with KIR-blocking; block-sensitive, detected only without KIR-blocking; or block-only, detected only with KIR-blocking. Cells without an assigned clonotype were excluded. Statistical comparisons were performed using two-sided Wilcoxon rank-sum tests with Benjamini–Hochberg adjustment for multiple testing. Only significant comparisons are indicated. **A)** KIR RNA module score across clonotype classes. Shared vs block-sensitive, adjusted p<0.001; block-sensitive vs block-only, adjusted p<0.001. **B)** Mean total KIR surface protein expression, calculated as the mean CITE-seq expression of CD158 (KIR2DL1/DS1/DS3/DS5), CD158b (KIR2DL2/DL3), and CD158e1 (KIR3DL1). Shared vs block-sensitive, adjusted p<0.001; block-sensitive vs block-only, adjusted p<0.001. **C)** Heatmap of KIR gene expression scaled per gene and shown as Z-scores. Statistical comparisons were performed using the unscaled RNA expression data. Significant differences were observed for KIR2DL1, KIR3DL1, and KIR3DL2. Shared vs block-sensitive: KIR2DL1, KIR3DL1, and KIR3DL2, adjusted p<0.001. Block-sensitive vs block-only: KIR2DL1, adjusted p<0.001; KIR3DL1, adjusted p=0.0033; and KIR3DL2, adjusted p=0.0038. **D)** Individual KIR surface protein expression measured by CITE-seq. KIR2DL1/DS1/DS3/DS5: shared vs block-sensitive, adjusted p<0.001; block-sensitive vs block-only, adjusted p<0.001. KIR2DL2/DL3: shared vs block-sensitive, adjusted p=0.0109.

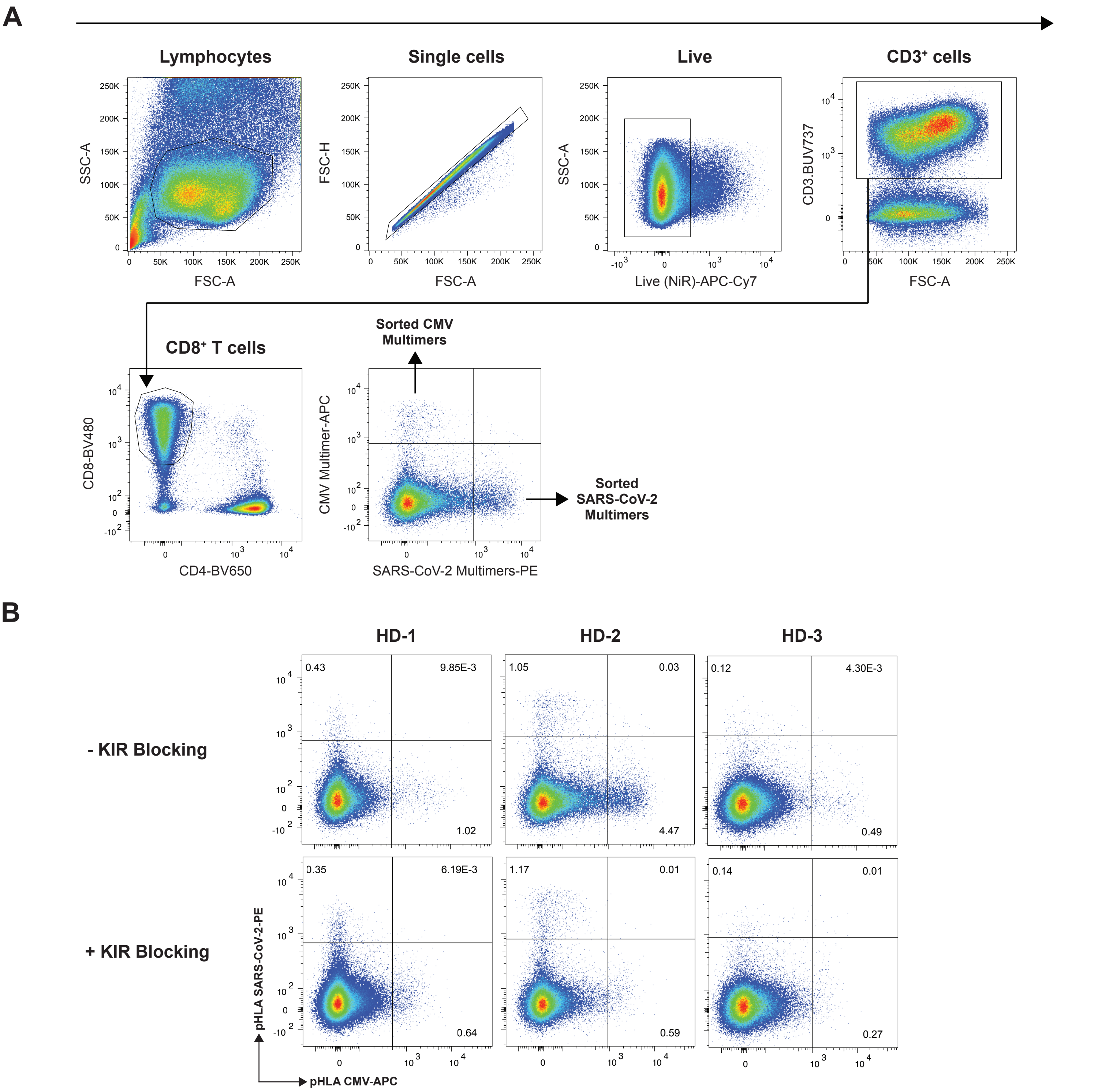


**Supplementary Figure 12: HLA-C*06:02 DNA-barcoded pHLA multimer analysis.**
**A)** Flow cytometry gating strategy used to identify and sort pHLA multimer⁺ CD8⁺ T-cells. **B)** Flow cytometry sorting plots showing HLA-C*06:02 pHLA multimer staining of CD8⁺ T-cells from three HLA-C*06:02-matched donors with or without KIR-blocking.

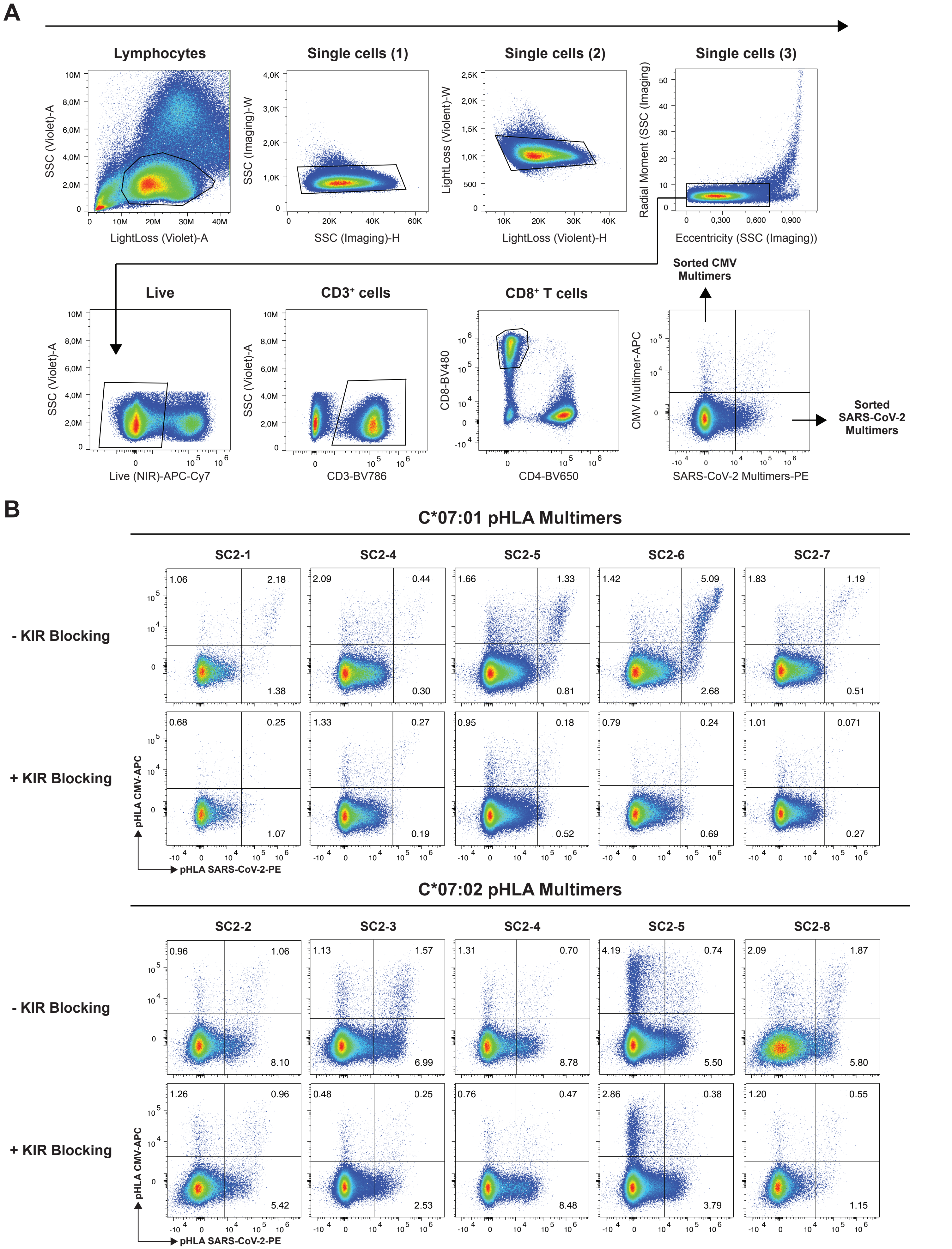


**Supplementary Figure 13: HLA-C1 DNA-barcoded pHLA multimer screening in SARS-CoV-2-exposed donors.**
**A)** Flow cytometry gating strategy used to identify and sort pHLA multimer⁺ CD8⁺ T-cells. SARS-CoV-2 pHLA multimers were labelled with PE-fluorophore, and CMV pHLA multimers were labelled with APC-fluorophore. **B)** Flow cytometry sorting plots showing pHLA multimer staining of CD8⁺ T-cells from HLA-C*07:01- and HLA-C*07:02-matched samples with and without KIR-blocking.

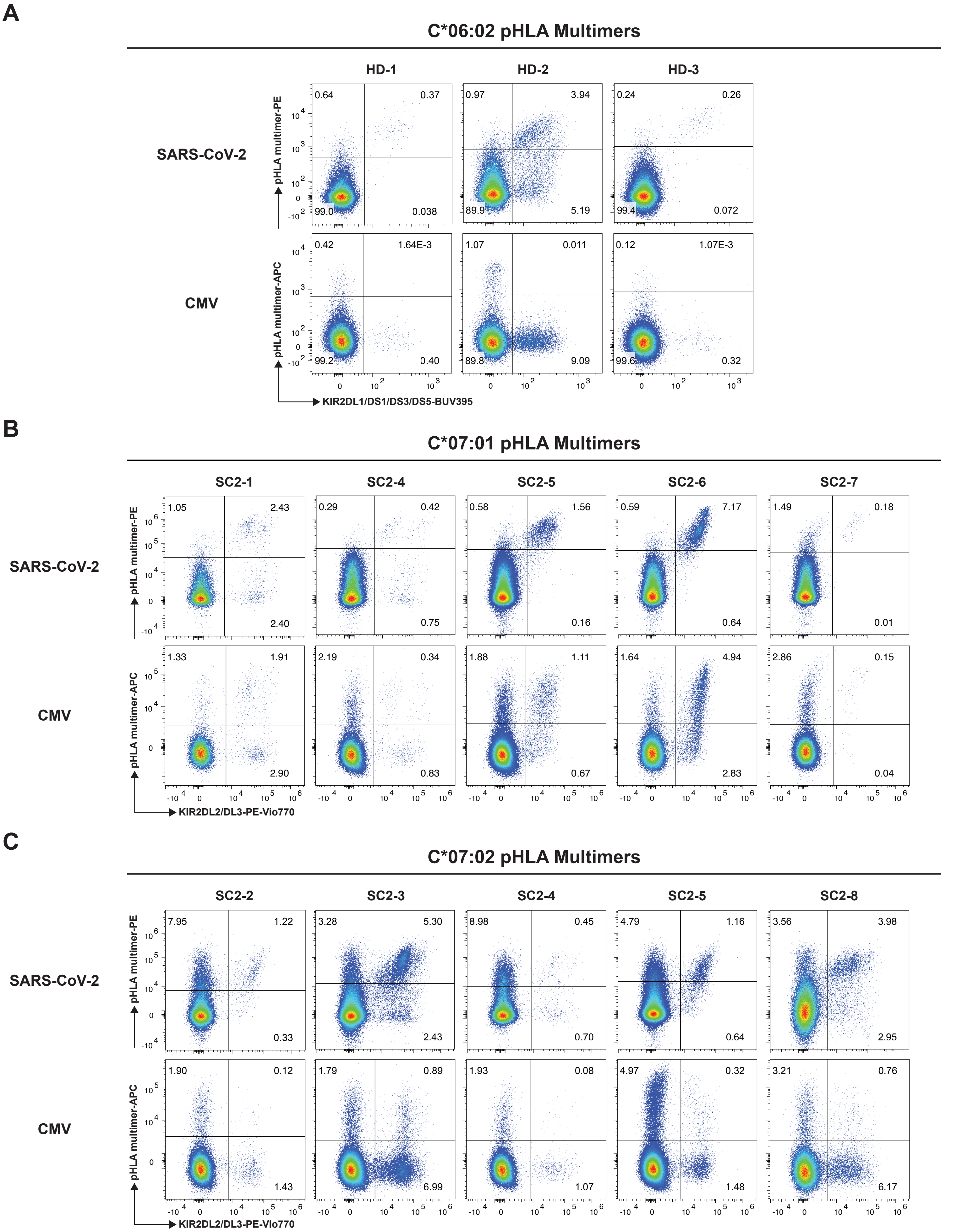


**Supplementary Figure 14: CMV- and SARS-CoV-2-derived HLA-C pHLA multimer staining and cognate KIR expression in DNA-barcoded screens.**
Flow cytometry plots showing HLA-C*06:02 pHLA multimer staining relative to KIR2DL1/DS1/DS3/DS5 **(A)**, HLA-C*07:01 pHLA multimer staining relative to KIR2DL2/DL3 **(B)**, and HLA-C*07:02 pHLA multimer staining relative to KIR2DL2/DL3 **(C)** expression in $CD8^+$ T-cells.

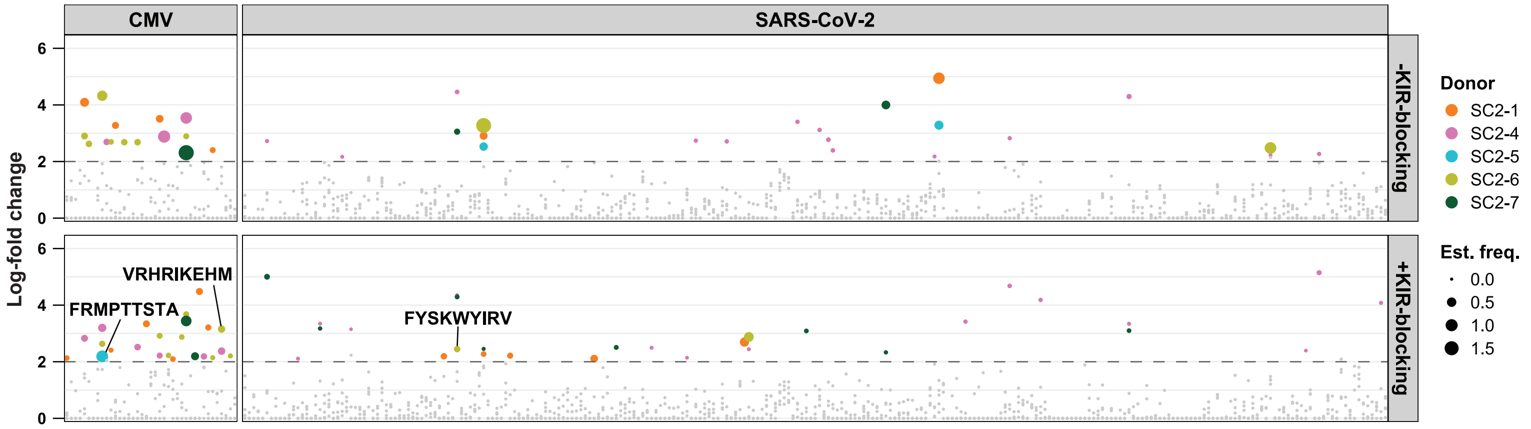


**Supplementary Figure 15: HLA-C*07:01 epitope detection with and without KIR-blocking and summary of DNA barcode-defined HLA-C epitope hits.**

DNA barcode-defined detection of HLA-C*07:01-restricted epitopes with and without KIR-blocking. HLA-C*07:01 is an HLA-C1 allotype. Significant responses were defined as Log2FC >2 and p<0.001. Colored markers indicate significant donor-specific responses, with marker size reflecting estimated frequency among CD8⁺ T-cells. Grey markers indicate non-significant enrichment. Selected epitopes are annotated.

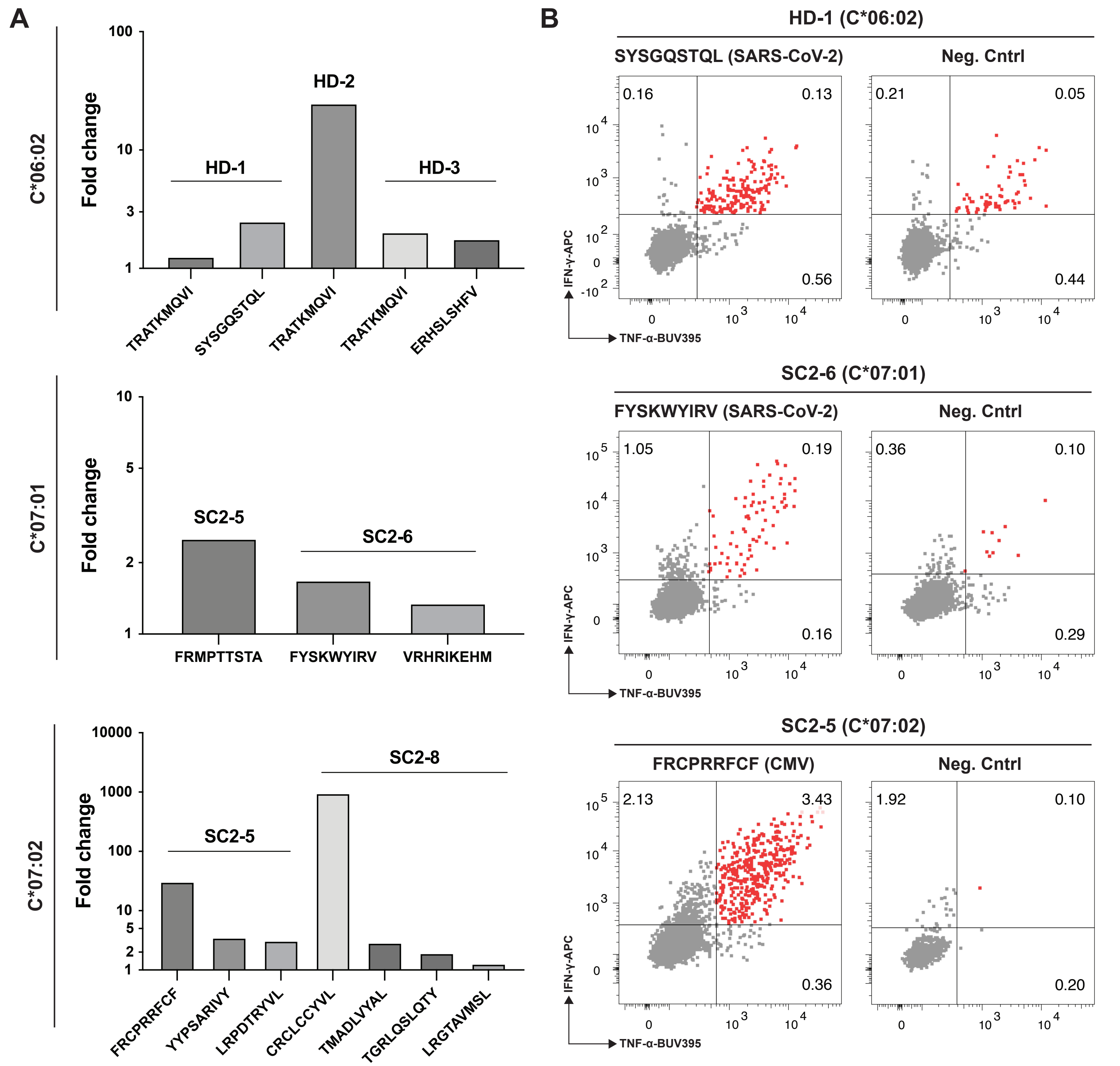


**Supplementary Figure 16: Functional validation of HLA-C-restricted epitopes identified using DNA-barcode based large-scale pHLA multimer analysis.**

PBMCs were expanded with individual HLA-C-restricted epitopes for 13 days before restimulation with the corresponding peptide. **A)** Peptide-induced IFN-γ and TNF-α production shown as fold change relative to the matched negative control and grouped by HLA restrictionand donor. Fold changes are displayed on a Log10 scale. **B)** Flow cytometry plots showing IFN-γ and TNF-α production following restimulation with a selected HLA-C*06:02-, HLA-C*07:01-, and HLA-C*07:02-restricted epitope respectively and their corresponding negative controls.
