## Supplementary Tables for "KIR Expression Defines a Transcriptionally Distinct CD8+ T-cell Population that Confounds Antigen-Specific T-cell Detection"

**Supplementary Table 1. HLA-A-, HLA-B- and HLA-C-restricted T-cell responses detected across RCC, NCSCLC, melanoma, and bladder cancer cohort.** p values compare the proportion of detected responses for each HLA restriction with the most frequent restriction within each cohort.

**Supplementary Table 2. MDS patient-specific neopeptide pHLA multimer screening.**
DNA-barcoded pHLA multimer screening results for neopeptides derived from an index MDS patient, including HLA-C*02:02-predicted neopeptides and additional HLA-A- and HLA-B-restricted neopeptide multimers included for comparison. For each peptide-HLA library, the table shows the mutated peptide sequence, amino acid change, barcode enrichment Log2FC, and associated p value in the MDS patient, one HLA-C*07:02-matched healthy control, and one HLA-C-mismatched healthy control.

**Supplementary Table 3. CMV- and EBV-derived HLA-C*02:02 peptide library used for combinatorial pHLA multimer staining.**
List of 46 CMV- and EBV-derived peptides predicted to bind HLA-C*02:02 and analyzed by combinatorial pHLA multimer staining. The table includes peptide sequence, viral origin, source protein, amino acid position, and predicted NetMHCpan binding rank.

**Supplementary Table 4: Viral HLA-C peptide libraries used for DNA-barcoded pHLA multimer screening.**
Peptide-HLA combinations are provided in separate sheets for HLA-C*06:02, HLA-C*07:01, and HLA-C*07:02. Each sheet includes viral origin, protein source, peptide sequence, peptide length, and predicted NetMHCpan 4.1 binding rank.

**Supplementary Table 4. Complete results from DNA-barcoded pHLA-C multimer screening with and without KIR-blocking.**
Complete screening results for 573 unique SARS-CoV-2- and CMV-derived peptides predicted to bind HLA-C*06:02 (n=287 peptides; n=3 donors), HLA-C*07:01 (n=298 peptides; n=5 donors), or HLA-C*07:02 (n=388 peptides; n=5 donors). Samples were analyzed in parallel with and without KIR-blocking. The table includes sample identifier, cohort, KIR-blocking condition, barcode count, Log2 fold change, p value, HLA allele, source protein, peptide sequence, viral origin, response classification, and estimated frequency. For the KIR-blocking variable, 0 indicates no KIR-blocking and 1 indicates KIR-blocking.

**Supplementary Table 6. Viral HLA-C peptide-specific binding events identified by DNA-barcoded pHLA multimer screening.**
Identified SARS-CoV-2- and CMV-derived HLA-C peptide-specific binding events from the DNA-barcoded pHLA multimer screens. The table includes HLA allele, viral origin, peptide sequence, protein source, number of donors with detectable binding, prevalence among HLA-

matched buffycoats, intracellular cytokine staining validation status, and previous reporting status.

**Supplementary Table 7. Streptavidin conjugates used for pMHC tetramer generation.**

**Supplementary Table 8. Antibody panels. For antibodies with multiple versions listed, any listed version may have been used.**

**Supplementary Table 9. Summary of titration results for fluorescence-labeled and purified KIR-blocking antibodies.**
